# DPP4 inhibition affects metabolism and inflammation associated pathways in hiPSC-derived steatotic HLCs

**DOI:** 10.1101/2024.11.06.622222

**Authors:** Christiane Loerch, Wasco Wruck, Julian Reiss, James Adjaye, Nina Graffmann

**Affiliations:** Institute for Stem Cell Research and Regenerative Medicine, Medical Faculty and University Hospital Düsseldorf, Heinrich Heine University, Düsseldorf, Germany; University College London, EGA Institute for Women’s Health, Zayed Center for Research into Rare Diseases in Children (ZGR), London, United Kingdom

**Keywords:** human induced pluripotent stem cells, Hepatocyte-like cells, Dipeptidyl peptidase 4 (DPP4), Cluster of Differentiation 26 (CD26), MAFLD, MASLD, NAFLD, Steatosis, Diabetes, Vildagliptin

## Abstract

**Background:** Metabolic dysfunction-associated steatotic liver disease (MAFLD) has a high prevalence and high co-morbidity for other diseases. Due to the complexity of this multifactorial disease, therapy options are still rather limited. We employed an *in vitro* pluripotent stem cell-based model to decipher potential disease-associated molecular pathways and to study the mode of action of prospective drugs. Dipeptidyl peptidase 4 (DPP4) or Cluster of differentiation 26 (CD26) is involved in inflammation, infections, immune disorders, type 2 diabetes, kidney disease and cancer.

**Methods:** We induced the steatosis phenotype in human induced pluripotent stem cell (iPSC) derived hepatocyte-like cells (HLCs) by oleic acid (OA)-feeding and confirmed regulation of clinically relevant pathways by NGS-based global transcriptomic analyses. Analysis of the secretome of steatotic HLCs revealed DPP4 as a potential key mediator of the disease. To further elucidate its role in the development of MAFLD, we inhibited DPP4 activity with Vildagliptin (VILDA) and analyzed the global transcriptome changes as well as specific gene and protein expression of steatosis-associated genes with and without DPP4 inhibition.

**Results:** MAFLD-associated pathways such as PPAR- and TNF signaling were differentially regulated in hiPSC-derived steatotic HLCs. We found increased hepatic DPP4 activity and secretion upon OA. Gene expression of fatty acid and purine metabolism and inflammation-associated pathways were regulated upon DPP4 inhibition.

**Conclusions:** Our HLC-model confirmed association of DPP4 with metabolism and inflammation which foster the development of MAFLD. Inhibiting DPP4 with VILDA partially relieved the steatotic phenotype on a global transcriptomic level.

**Impact and implications:** Given the difficulties of identifying suitable anti-MAFLD drugs, novel model systems are urgently needed. Our *in vitro* HLC-model reproduced DPP4-dependent aspects of the disease and responded positively to Vildagliptin treatment. Further elucidation of the role of DPP4 in the etiology of MAFLD and other diseases is warranted.

## Background

Steatotic liver diseases are an increasing health burden for industrialized countries all over the globe(1–4). Early, reversible stages involve steatosis and steatohepatitis, resulting in fibrosis, while nascent cirrhosis and hepatocellular carcinoma are non-reversible and life-threatening. Primary causes for steatosis are elevated fatty acid flux from adipose tissue, a high-fat diet, or elevated blood glucose levels. Thus, obesity and Type 2 diabetes mellitus (T2DM) are direct diagnostic criteria for Metabolic dysfunction-associated fatty/steatotic liver disease (MAFLD/MASLD)(2). MAFLD is a complex, multisystem disease, with serious implications on the whole body, including cardiovascular disease and chronic kidney diseases(5, 6). Moreover, its multifactorial character and tissue heterogeneity is not only found in the clinic but is also shown in the molecular response to steatosis(7–9). These factors complicate research and to date resmetirom is the only FDA approved drug for treating MASLD(6, 10, 11).

Dipeptidyl peptidase 4 (DPP4) is a serine protease with catalytic activity for various substrates(12). The most prominent ones are the incretins-Glucagon like peptide 1 (GLP-1) and Glucose-dependent insulinotropic polypeptide (GIP1). However, DPP4 not only interferes with incretin signaling, but also cleaves chemotactic peptides with implications for the inflammatory response(12, 13). It is therefore considered a hepatokine, upregulated in metabolic liver disease and a driver of inflammation(13). Furthermore, DPP4 is known to be involved in various etiologies, such as immune disorders, fibrosis and cancer(14). Gliptins -potent DPP4 inhibitors-are used for the treatment of T2DM, prolonging the postprandial incretin answer(13, 15–17). Vildagliptin (VILDA) shows additional protective effects on hepatocytes by reducing hepatic triglyceride load as well as aminotransferase levels(18). Since the inflammatory response was reduced, it might interfere with the traits towards steatohepatitis, thus indicating a beneficial effect on the progression of the disease(19, 20).

In this study, we differentiated human patient-derived induced pluripotent stem cells (iPSCs) to hepatocyte-like cells (HLCs) by using our previously published, efficient 2D differentiation protocol(21). We induced the steatosis phenotype and provide insights into the hepatocyte-specific contribution to the disease and the potential role of DPP4 for the interplay between metabolism and inflammation.

## Methods

### Human induced pluripotent stem cell (iPSC) culture and hepatocyte-like cell (HLC) differentiation

The use of iPSC lines for this study was approved by the ethics committee of the medical faculty of Heinrich-Heine University under the numbers 5013 and 5704. iPSCs were cultivated on Matrigel (Corning) coated 6-well plates with daily changes of StemMACs medium (Miltenyi). Once iPSCs attained 90% confluency, they were colony-split by addition of PBS w/o magnesium or calcium (PBS-/-) (Gibco) and incubated for approximately 3 min at room temperature (RT). Colonies were detached from the surface with a cell scraper and centrifuged at 40 *xg* for 3 min. The pellet was carefully resuspended and clumps of colonies were seeded in a ratio of 1:6.

iPSCs were differentiated according to our previously published protocol(21). In brief, 1.04 x10^5^ iPSCs/ cm^2^ were seeded onto Matrigel-coated dishes. Definitive endoderm (DE) was induced by 1-3 days of 2.5 µM CHIR99021 (Stemgent) and 3-5 days of 100 ng/ml Activin A (Peprotech) in RPMI medium. Hepatic endoderm (HE) medium was fed for 4 days, and 1% DMSO was added with medium changes every day. HLC medium was fed for 12-15 days, with medium changes every other day. 1 µM insulin (Sigma-Aldrich), 10 ng/mL hepatocyte growth factor (HGF) (Peprotech), 25 ng/ml dexamethasone (Dex) (Sigma-Aldrich), and 20 ng/mL recombinant human Oncostatin M (rhOSM209a.a) (Immunotools) were freshly added to the medium.

### Immunocytochemistry

Cells were washed with PBS-/-, fixed with 4% PFA for 10 min at RT, and washed 3x with PBS-/-. For intracellular staining, the cells were permeabilized with 0.5% Triton-X-100 (Sigma-Aldrich) in PBS-/-for 10 min at RT, and blocked with 3% BSA in PBS-/-for 1 h at RT. After incubation with primary antibodies (Supplementary material) at respective dilutions (table S2) overnight at 4°C, unbound antibodies were washed off 3x with PBS-/-. Secondary antibodies against the respective host IgG were incubated for 1 h at RT and washed 3x with PBS-/-. (Confocal) microscopy was performed, using a LSM 700 microscope (Zeiss) and images were processed with ZEN software (Zeiss).

### Quantitative reverse transcription PCR (qRT-PCR)

RNA was isolated using the Direct-zol RNA isolation kit (Zymo Research), following the manufacturer’s instructions. 500 ng of RNA was reverse transcribed to cDNA using the TaqMan reverse transcription kit (Life technologies). qRT-PCR was performed using the VIIA7 machine and the power SYBR green master mix (all Life technologies). Expression of mRNA was presented as log2-fold-change in comparison to the housekeeping gene and the control condition (primer sequences are found in the supplementary material table S1).

### Western Blot

Cells were lysed in RIPA buffer containing protease and phosphatase inhibitors (all Sigma-Aldrich). 15 - 30 µg proteins were separated on NuPAGE 4 to 12% Bis-Tris protein gels (Life Technologies) and wet-blotted onto 0.45 µm nitrocellulose membranes (Amersham). After blocking with 5% non-fat milk (ROTH) in TBS-T buffer, the membranes were incubated with the respective primary antibodies (Supplementary material, table S2) overnight at 4 °C. After 3x washing with TBS-T buffer, fluorescence labeled secondary antibodies (Licor) against the host IgG were incubated for 1 h at RT at a 1:10000 dilution. Unbound antibodies were washed off with TBS-T buffer and the fluorescence signal was detected at 680 nm and 800 nm by using the ChemiDoc MP Imaging system (Bio-Rad). Quantification was performed using the Image Lab 6.0.1 software with lane background subtraction using disk size 1.

### Cytochrome P450 activity measurement

P450-Glo^TM^ CYP3A4 and CYP2D6 assays (Promega) were used to measure Cytochrome P450 activity. Cells were incubated with 3 µM Luciferin-IPA or 10 µM Luciferin-ME EGE, respectively, in William’s E Medium (Gibco) for 1 h at 37 °C. After incubation with the detection reagent, luminescence was measured in technical triplicates with a luminometer (Lumat LB 9507, Berthold Technologies).

### Steatosis induction by OA-feeding and Vildagliptin (VILDA) treatment

Oleic acid (Calbiochem) was bound to 14% (w/v) fatty acid free BSA (ROTH) in 0.1 M TRIS pH 8.0 for 1 h at 37 °C, and stored at 4 °C. After testing different concentrations and time periods of OA treatment, we selected 400 µM OA for 7 days to induce steatosis. From day 15-17 of HLC differentiation on, the cells were fed with complete HLC medium, supplemented with 400 µM OA or the respective volume of TRIS-BSA as mock-treatment. The medium was changed every other day for 7 days. A final concentration of 30µM VILDA (Sigma-Aldrich) dissolved in DMSO was fed to the cells after 48 h OA-/mock-induction for 5 days with medium changes every other day.

### Next generation sequencing and analysis of deep sequencing data

3’RNA-Seq was performed on a NextSeq2000 sequencing system (Ilumina) at the core facility Biomedizinisches Forschungszentrum Genomics and Transcriptomics laboratory (BMFZ-GTL) of Heinrich-Heine University Duesseldorf. The HISAT2 (version 2.1.0) software(26) was employed to align the fastq files received from the BMFZ-GTL core facility against the GRCh38 genome. For detailed description of the integration of the data please refer to the supplementary material, methods section.

### GO and pathway analysis

Subsets of genes expressed exclusively in one condition in the Venn diagram analysis and up- and down-regulated genes according to the criteria for differentially expressed genes (limma test, p-value <0.05 and fold change >1.5) (were subjected to over-representation analysis of gene ontologies (GOs) and KEGG (Kyoto Encyclopedia of Genes and Genomes) pathways(27). The hypergeometric test built-in in the R base package was used for over-representation analysis of KEGG pathways, which had been downloaded from the KEGG database in February 2023. The GOstats R package(28) was employed to determine over-represented GO terms. The most significant GO terms and KEGG pathways were displayed in dotplots via the R package ggplot2(29).

### Enzyme-linked Immunosorbent assay (ELISA)

Secreted DPP4 was detected from supernatants 48 h after feeding using the human DPP4/CD26 DuoSet ELISA (R&D Systems) as described by the manufacturer. Optical density was measured using the EPOCH2 spectrophotometer (BioTek) at 450 nm with wavelength correction at 540 nm. 4-PL curve fitting was performed to calculate concentrations.

### Enzyme activity assay

DPP4 activity was measured in OA-/ VILDA-treated HLCs, using the Dipeptidyl peptidase IV (DPP4) Activity Assay Kit (Fluorometric) (Abcam), following the manufacturer’s instructions. Fluorescence signal was measured on a Spectrophotometer (Tecan) at Ex/Em = 360/460 nm.

### Cytokine array

Supernatants of three biological replicates were harvested 24 h after medium change and Proteome Profiler Human XL Cytokine array (R&D Systems) analysis was performed following the manufacturer’s protocol and signals were detected, using the Fusion FX instrument (PeqLab). Analysis and quantification was performed using the FIJI/ImageJ software(30) and the Microarray Profile plugin by Bob Dougherty and Wayne Rasband (https://www.optinav.info/MicroArray_Profile.htm, accessed on 21 December 2022). For details of the image analysis and follow-up normalization in the R/Bioconductor environment(31) we refer to the description in our previous publication(32). Cytokines were considered differentially expressed satisfying the criteria: detection p-value < 0.05 in at least one condition, fold change > 1.2 and limma-p-value < 0.05. The function heatmap.2 from the gplots package(33) and the R-builtin function barplot were applied for heatmap and bar plots.

### Statistics

Student’s unpaired two-tailed t-test or ordinary one- or two-way ANOVA followed by Tukey’s, Dunnet’s, or Sidak’s multiple comparison test were conducted to calculate significances using GraphPad Prism 8.0.2 software.

## Results

### iPSCs-derived hepatocyte-like cell (HLC) differentiation from four individuals

Induced pluripotent stem cells (iPSCs) derived from four individuals, two healthy controls (Cntrl 1, Cntrl 2)(22, 23) and two steatosis patients (Stea 1, Stea 2)(24, 25)(Table 1), were differentiated into HLCs following our recently published protocol(21). Representative immunocytochemistry of Cntrl 2 shows the expression of Octamer binding transcription factor 4 (OCT4) in iPSCs, SRY-box transcription factor 17 (SOX17) in definitive endoderm (DE), and Alpha-fetoprotein (AFP) in hepatic endoderm (HE) (Fig. 1A). HLCs were stained for the epithelial marker E-Cadherin (E-CAD), Hepatocyte nuclear factor 4 alpha (HNF4a) and Albumin (ALB) (Fig.1A). Representative pictures of the other cell lines are provided in the supplementary material section (Fig. S1 and S2A-C). HLCs showed significant increase of gene expression of the HLC-markers *ALB* and Cytochrome P450 family members *CYP3A4* and *CYP2D6* in comparison to the iPSC-stage (Fig. 1B). Representative gene expression of *OCT4* and *SOX17* in DE, as well as *AFP* in HE stage is provided in Fig. S2D. Protein expression of AFP and ALB is shown in comparison to the housekeeping protein beta-Actin (bActin) in HLCs derived from Cntrl 1 and Cntrl 2 (Fig. 1C). HLCs’ functionality was confirmed by measuring CYP3A4 and CYP2D6 activity for the four cell lines (Fig. 1D).

**Table 1:**
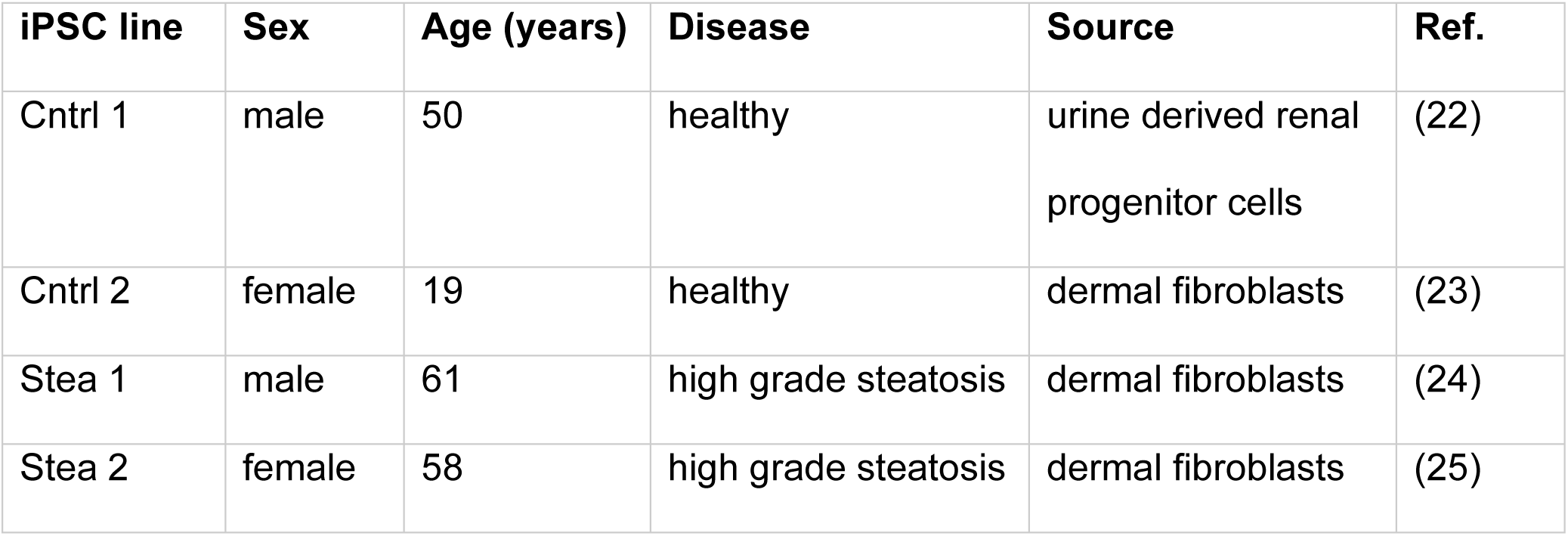
iPSC lines used in this study.

| iPSC line | Sex | Age (years) | Disease | Source | Ref. |
| --- | --- | --- | --- | --- | --- |
| Cntrl 1 | male | 50 | healthy | urine derived renal progenitor cells | (22) |
| Cntrl 2 | female | 19 | healthy | dermal fibroblasts | (23) |
| Stea 1 | male | 61 | high grade steatosis | dermal fibroblasts | (24) |
| Stea 2 | female | 58 | high grade steatosis | dermal fibroblasts | (25) |

**Fig. 1.**
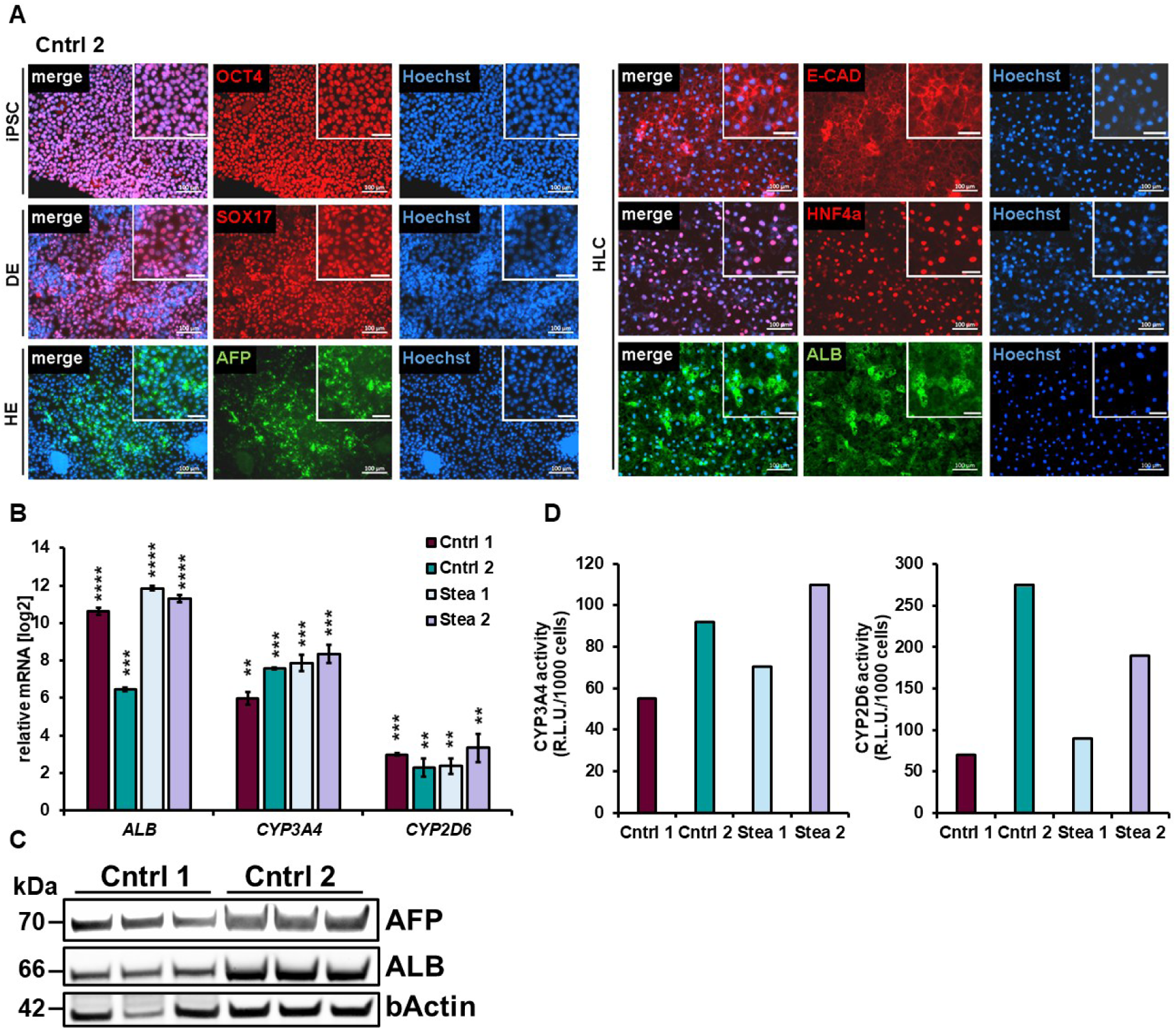
Characterization of HLCs. **(A)** Representative immunocytochemistry of cell line Cntrl 2 during differentiation showing respective markers OCT4 in iPSCs, SOX17 in definitive endoderm (DE), and AFP in hepatic endoderm (HE). For hepatocyte-like cells (HLCs), the epithelial marker E-CAD, HNF4a and ALB are shown. Scale bars = 100 µm. **(B)** Gene expression of *ALB* and Cytochrome P450 family members *CYP3A4* and *CYP2D6* in HLCs (n=3) derived from four cell lines in comparison to iPSC-stage (n=2), shown in means of two or three biological replicates ±SE respectively, normalized to the housekeeping gene *RPLP0*. Two-tailed unpaired students’ T-test was performed to calculate significances (*p < 0.05, **p < 0.01, ***p < 0.001). **(C)** Cropped WB of AFP, ALB and bActin in HLCs derived from Cntrl 1 and Cntrl 2 (n=3). Uncropped full-length blots can be found in the supplementary Fig. S9. **(D)** Representative cytochrome P450 activity of HLCs is for CYP3A4 and CYP2D6 in relative light units (R.L.U.) per 1000 cells in technical triplicates (n=3).

### Oleic acid (OA) induces steatosis phenotype in HLCs

To induce the steatosis phenotype in iPSC-derived HLCs, we treated HLCs of all four cell lines on day 15-17 of differentiation with 400 µM oleic acid (OA) for 7 days. After OA induction, we detected the formation of Perilipin-2 (PLIN2) coated lipid droplets by immunocytochemistry in all cell lines (Fig. 2A/B and Fig. S3). Interestingly, from a visual impression, it seemed that Cntrl 1 showed less lipid droplets than Cntrl 2, indicating a cell line specific difference in the build-up of lipid droplets. In accordance with previous findings, we did not detect a disease specific difference in the lipid load w or wo OA, but rather a cell line specific effect(7). Previous findings indicated distinct gene expression profiles in response to OA-treatment related to the steatosis background of HLCs. To put these observations in perspective, we analyzed the global transcriptomic changes upon OA induction.

**Fig. 2.**
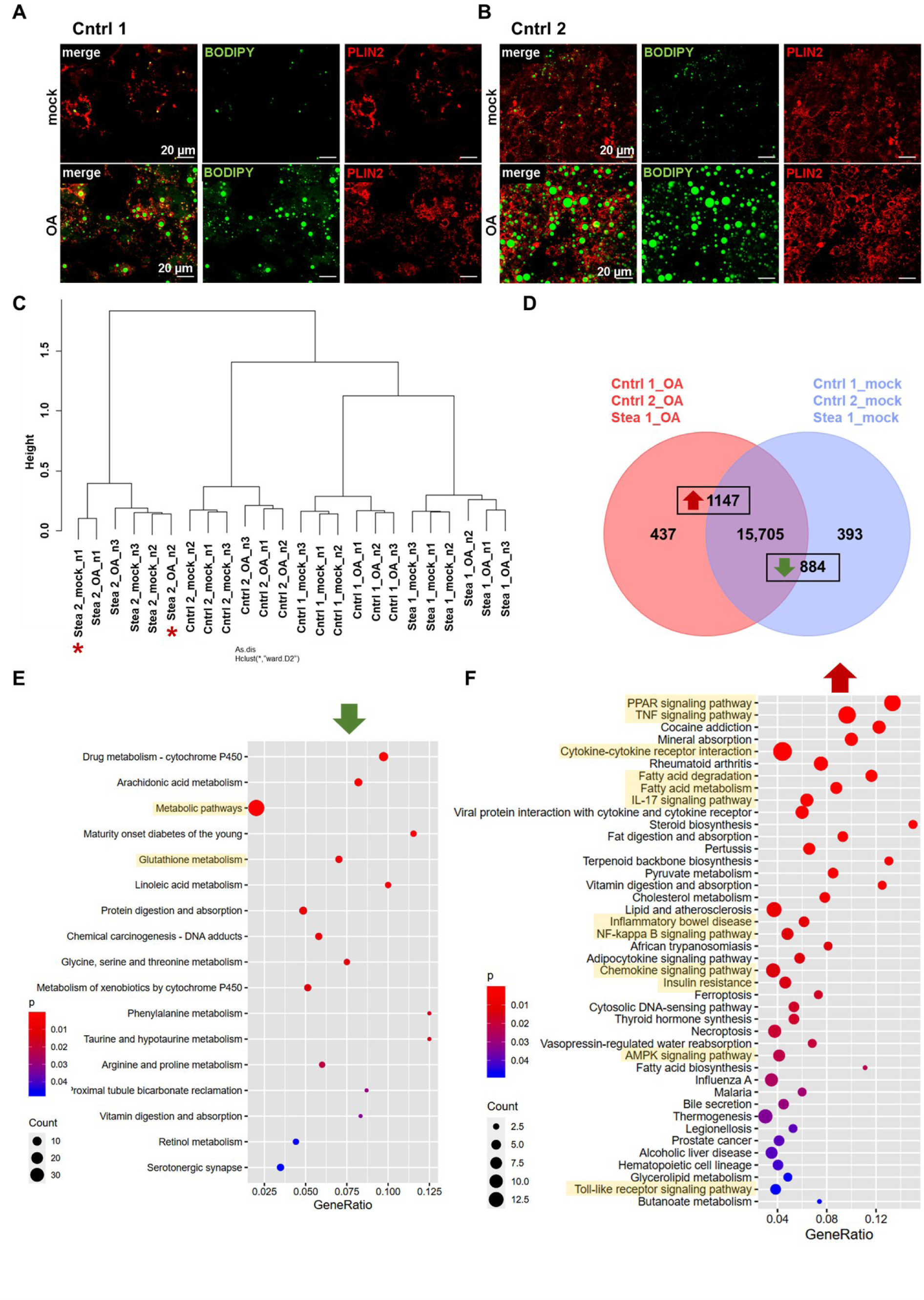
Confirmation of steatosis phenotype in HLCs. **(A/B)** Representative immunofluorescence and BODIPY 493/503 staining of HLCs derived from Cntrl 1 and Cntrl 2 treated with 400 µM OA and respective control (mock) for 7 days, PLIN2 is shown in red, fatty acids are stained in green. Scale bars = 20 µm. **(C)** Hierarchical cluster dendrogram of global transcriptomic changes upon OA treatment of HLCs derived from cell lines Cntrl 1, Cntrl 2, Stea 1, Stea 2 in three biological replicates (n=3). Ambiguous clustering of gene expression in Stea 2 is marked by red asterisks (correlation in supplementary material table 1). **(D)** Venn diagram of gene expression from OA treated HLCs derived from cell lines Cntrl 1, Cntrl 2, Stea 1 indicating 437 solely expressed genes upon OA, and 393 solely expressed genes upon mock treatment and 15,705 genes expressed in common. Among exclusively and commonly expressed genes 1,147 and 834 genes were significantly up- and down-regulated, respectively. **(E/F)** Dot plots of KEGG-associated pathway analysis of significantly down **(E)**- and up **(F)**-regulated genes (gene lists in supplementary table S3).

RNA-seq was performed and revealed gene expression clustering according to the treatment and based on the genetic background of each cell line (Fig. 2C). However, two samples of HLCs derived from cell line Stea 2, clustered ambiguously, (Fig. 2C, red asterisks). To prevent biases due to the genetic background of this cell line, we excluded Stea 2 in subsequent analysis. We identified 15,705 genes expressed in common in both conditions in cell lines Cntrl 1, Cntrl 2 and Stea 1, while 437 genes were exclusively expressed under OA in comparison to 393 genes solely expressed genes under mock treatment. Combining the exclusively expressed genes and genes expressed in common, we found 1,147 genes significantly upregulated and 884 genes significantly downregulated (Fig. 2D) after OA treatment. KEGG-associated pathway analysis revealed, among others, genes of the glutathione pathway and metabolic pathways significantly downregulated upon OA-treatment throughout Cntrl 1, Cntrl 2 and Stea 1 cell lines (Fig. 2E). Confirming results from a previous study(7), we found genes associated with KEGG pathways of Peroxisome proliferator activated-receptor (PPAR)-, Adenosine monophosphate-activated protein kinase (AMPK)-signaling and fatty acid metabolism significantly upregulated (Fig. 2F). Furthermore, we detected genes belonging to inflammation-related pathways significantly upregulated upon OA, such as Tumor necrosis factor (TNF) signaling and NF-kappa-B pathway. Interestingly, also genes of the inflammatory bowel disease and insulin resistance pathways were upregulated upon OA (Fig. 2F). This KEGG-associated pathway analysis confirmed the induction of the steatosis phenotype by OA supplementation in the three cell lines Cntrl 1, Cntrl 2 and Stea 1. We performed Pearson’s correlation heatmap analysis of genes associated with relevant KEGG-pathways (Fig. 3A). Notably, this revealed a change of clustering according to the treatment and independent of the genotype, indicating the relevance of these genes for the phenotype. Taken together, global transcriptomic analyses of Cntrl1, Cntrl2, and Stea1 cell lines confirmed successful steatosis induction but did not reveal an altered susceptibility of the patient derived cells.

**Fig. 3.**
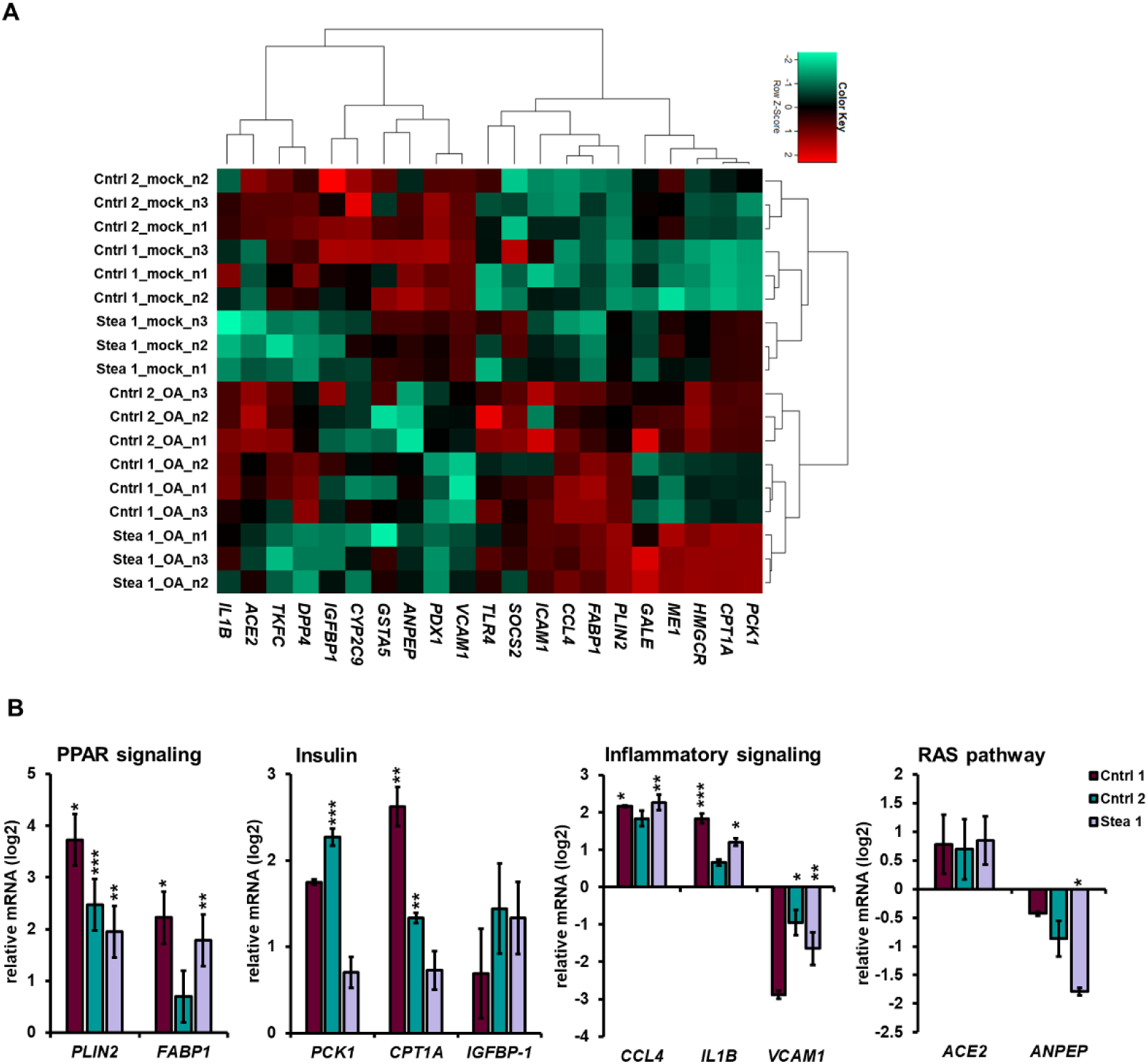
OA-treatment induces differential gene expression. **(A)** Pearson’s correlation heatmap analysis of the gene expression of members of OA-dysregulated pathways like fatty acid metabolism (*PLIN2*, *FABP1*), metabolic pathways/ glutathione metabolism (*ANPEP, GSTA5*) and insulin resistance (*PCK1*, *CPT1A*, *IGFBP1*, *SERPINA7*, *PDX1*). **(B)** qRT-PCR analysis of the mRNA expression of *PLIN2*, *FABP1, PCK1*, C*PT1A*, *IGFBP1*, *CCL4*, *IL1B, VCAM1 ACE2*, and *ANPEP* relative to mock-treated HLCs from cell lines Cntrl 1, Cntrl 2 and Stea 1 normalized to mock-treated HLCs derived from each cell line as means of two or three biological replicates ± SE (n=2 or n=3), normalized to the housekeeping gene *RPLP0*. Two-tailed unpaired Student’s T-test was performed to calculate significances (*p < 0.05, **p < 0.01, ***p < 0.001).

To confirm the global gene expression changes, we performed qRT-PCR for genes associated with relevant pathways (Fig. 3B). We found significant increases of PPAR-pathway associated genes, *PLIN2* and fatty acid binding protein-1 (*FABP1)* upon OA-treatment in at least 2 out of our 3 cell lines. Regarding insulin signaling associated genes, phosphoenolpyruvate carboxykinase-1 (*PCK1*) was significantly upregulated in Cntrl 2, and a non-significant trend towards upregulation was detectable in Cntrl 1 and Stea 1. Carnitine palmitoyltransferase-1A (*CPT1A*) was significantly upregulated in Cntrl 1 and 2. Interestingly, in contrast to the RNA-seq results, insulin-like growth factor binding protein-1 (*IGFBP1)* showed a tendency to upregulation upon OA, however not significantly. Considering genes associated with inflammation, a significant upregulation of CC-chemokine ligand 4 (*CCL4*) and interleukin 1beta (*IL1B)* expression was detected in Cntrl 1 and Stea 1, while in Cntrl 2 the increase was not significant. We found a significant reduction in the expression of vascular cell adhesion molecule-1 (*VCAM1*) in Cntrl 2 and Stea 1. Furthermore, we found members of the renin-angiotensin-system (RAS) differentially regulated in our model. For example, Angiotensin converting enzyme 2 (*ACE2*) tended to be upregulated in the RNA-seq data, a non-significant trend that could be confirmed by qRT-PCR. In addition, we detected a reduction of alanyl aminopeptidase (*ANPEP*), which is also involved in the glutathione metabolism (reactive oxygen species (ROS)-regulation), in all three cell lines, albeit only significantly in Stea 1. The tendency of up- and down-regulation of the qRT-PCR data in Fig. 3B confirms the direction of regulation in the RNAseq data. Together, these findings strengthen the validity of our model, because MAFLD-associated pathways were differentially regulated upon OA. They further underline the importance of the investigated genes as their differential expression was independent of the genetic background.

### Dipeptidyl peptidase 4 (DPP4) is secreted upon OA treatment

We analyzed the supernatant of OA-treated HLCs for released signaling proteins. Among others, we found a significant increase of DPP4, also known as Cluster of Differentiation 26 (CD26), upon OA-induction (Supp Fig. S4B/C). We confirmed this tendency of increase upon OA-treatment in Cntrl 1, Cntrl 2, and Stea 1 by ELISA (Fig. 4A) (0.95±0.11 ng/mL, 0.22±0.04 ng/mL, 2.51±0.27 ng/mL DPP4 under mock conditions and 3.94±0.38 ng/mL, 1.45±0.35 ng/mL, 5.53±0.02 ng/mL, under OA respectively).

**Fig. 4.**
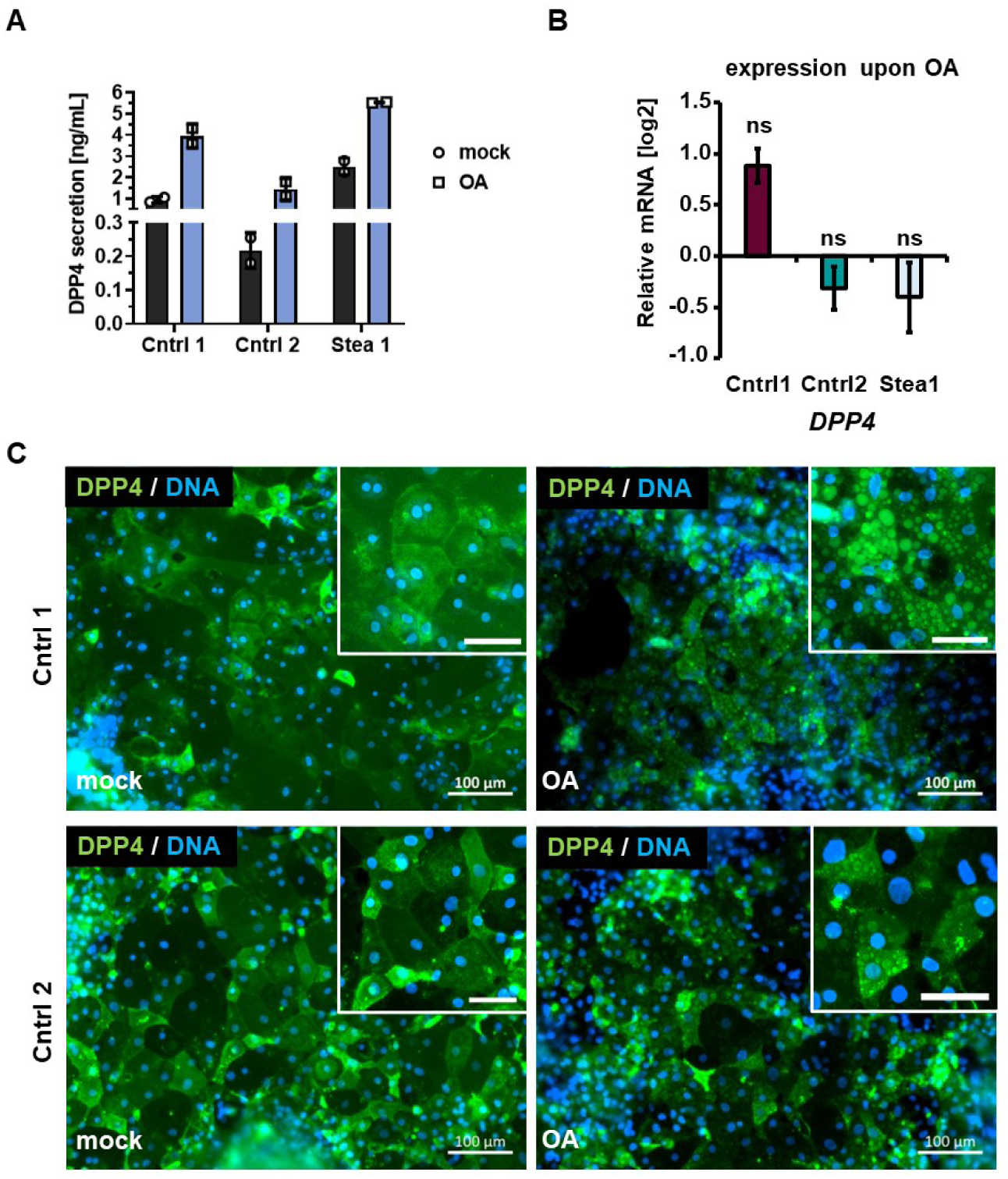
DPP4 is secreted upon OA-treatment. **(A)** Secretion of Dipeptidyl peptidase (DPP4) upon OA-treatment measured by ELISA-detection, showing secreted DPP4 as the mean of two biological replicates (n=2 ±SD). **(B)** Gene expression of *DPP4* upon OA treatment relative to mock treatment in all cell lines, shown are means of three biological replicates, normalized to the housekeeping gene *RPLP0* (n=3 ±SE). Two-tailed unpaired Student’s T-test was performed to calculate significances (ns p> 0.05) between OA and mock treatment for each cell line. **(C)** Representative immunocytochemistry of DPP4 under mock and OA conditions in cell lines Cntrl 1 and Cntrl 2. Scale bars represent 100 µm and 50 µm in the zoom-in.

Interestingly, there was a greater increase of DPP4 secretion upon OA in cell lines derived from healthy individuals, compared to patient derived HLCs. DPP4 levels increased approximately 4.5-to 5-fold in Cntrl 1 and 2 while Stea 1 showed a DPP4 increase of 2-fold respectively. This indicates that the control cell lines are able to increase DPP4 secretion more strongly in response to OA. To gain insights into the mechanisms underlying upregulation of DPP4 upon OA-treatment, we analyzed the gene expression in the three cell lines, however we could not detect a significant change upon OA-induction (Fig. 4B). To elucidate the role of DPP4 in steatosis, we focused for further analyses on the Cntrl cell lines, because of the stronger induction of DPP4 secretion upon OA. Similar to the gene expression of *DPP4*, we did not detect a prominent change in the protein localization or amount upon OA (Fig. 4C).

### Vildagliptin reduces DPP4 activity

Vildagliptin (VILDA) was tested in a phase-4 study (ID NCT01356381) to elucidate its potential use for treating steatosis patients(34). However, whether VILDA improves the hepatic phenotype directly or by incretin regulation, is yet to be elucidated. To shed light on its direct effects on hepatocytes, we induced the steatosis phenotype in our HLCs by a pre-treatment with OA for 48 h followed by incubation with 30 µM VILDA for a total of 5 days simultaneously with OA. We did not detect significant difference in *DPP4* expression on both the RNA and protein level upon the treatments in comparison to mock w/o VILDA (Fig. 5A-C), while we detected an expectedly strong increase for PLIN2 after OA induction in Cntrl 1 and 2 HLCs (Fig. 5B/C).

**Fig. 5.**
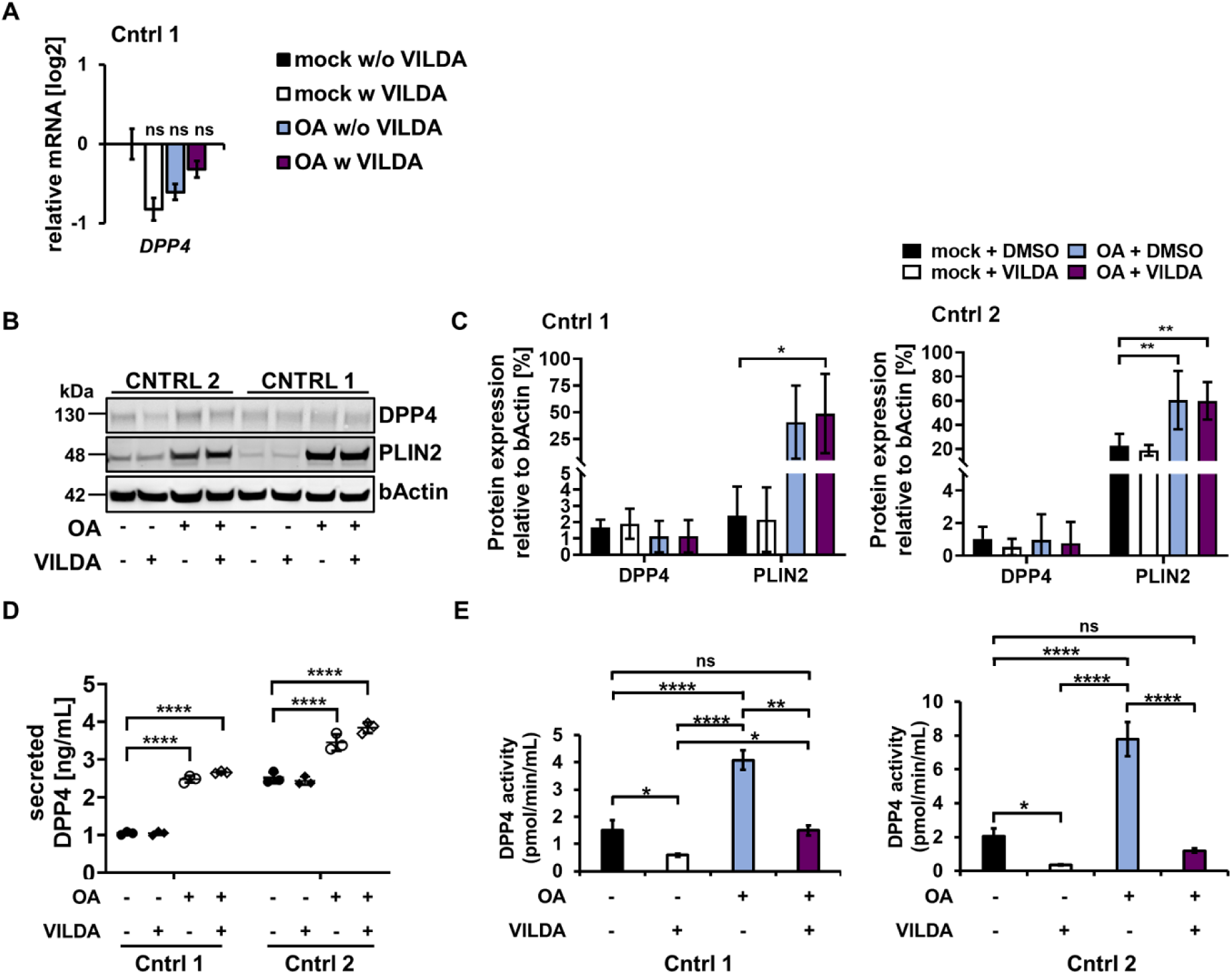
Effects of VILDA on OA-induced HLCs. **(A)** Gene expression of *DPP4* in Cntrl 1 HLCs upon OA-treatment with (w) and without (w/o) 30 µM Vildagliptin (VILDA) for 5 days, in comparison to mock w/o VILDA, shown are means of three biological replicates, normalized to the housekeeping gene *RPLP0* (n=3 ±SE). Ordinary one-way ANOVA followed by Tukey’s multiple comparisons was performed to calculate significances (ns p> 0.05). **(B)** Representative cropped WB of DPP4 and PLIN2 upon OA-treatment w and w/o VILDA. Uncropped full-length blots can be found in the supplementary Fig. S9. **(C)** Protein expression relative to bActin in comparison to mock w/o VILDA. Shown are means of three biological replicates (n=3 ± SD). Ordinary two-way ANOVA, followed by Tukey’s multiple comparison test was performed to calculate significances (*p < 0.05, **p < 0.01). Uncropped full-length blots can be found in the supplementary Fig. S9. **(D)** DPP4 secretion upon OA-treatment w and w/o VILDA. Shown are means of three biological replicates (n=3 ±SD). Ordinary two-way ANOVA, followed by Dunnet’s multiple comparison test was performed to calculate significances (**p < 0.01, ***p < 0.001, ****p < 0.001). **(E)** DPP4 activity in HLCs upon OA-treatment w and w/o VILDA was analyzed by measuring the proteolytic activity over time. Ordinary one-way ANOVA and Tukey’s multiple comparison test was performed to calculate significances (*p < 0.05, ****p < 0.001).

Considering the secretion of DPP4, we confirmed the previously detected increase of DPP4 upon OA w and w/o VILDA for both cell lines in comparison to the mock treatment. However, no significant difference upon VILDA treatment was detectable (Fig. 5D). Nevertheless, we detected a significant increase of DPP4 activity for both cell lines upon OA treatment which was significantly reduced when HLCs were treated with OA and VILDA together (Fig. 5 E). VILDA is capable of reducing DPP4 activity to the level detected under mock conditions, with no significant difference between OA w VILDA and mock w/o VILDA. These findings confirm that VILDA mainly acts on DPP4 activity, and neither on its gene or protein expression, nor on its secretion.

### Inhibition of DPP4 activity might reduce inflammatory progression leading to the disease phenotype

To test whether VILDA further affects other genes involved in the steatosis phenotype, we analyzed the global gene expression upon OA w and w/o VILDA in cell line Cntrl 1. To gain first insights on potential effects of DPP4 inhibition, we selected Cntrl1 for in depth analysis by NGS which will give directions for the necessary follow up studies with other cell lines. First level dendrogram analysis revealed clustering according to mock- and OA-treatment, however not according to the VILDA treatment, indicating a stronger impact of OA on the gene expression in comparison to VILDA (Fig. S5A). To confirm the previously detected gene expression pattern in response to OA, we performed KEGG-associated pathway analysis of the gene expression upon OA w/o VILDA in comparison to mock w/o VILDA (Fig. S5B/C). Indeed, we found similar pathways to be differentially regulated upon OA w/o VILDA in comparison to the previous data, indicating that the solvent reagent had no impact on the OA-response (Fig. S5B/C).

In addition to the 14,118 genes expressed in common in all four conditions, we found exclusively expressed gene sets for every condition as indicated by Venn analysis (Fig. 6A). We found 91 and 339 exclusively expressed genes upon OA w VILDA and mock w VILDA, respectively. 62 and 115 genes were exclusively expressed upon OA w/o VILDA and mock w/o VILDA treatment, respectively. KEGG- and Gene Ontologies (GO)-associated pathway analyses for the exclusively expressed genes upon the different conditions did not reveal characteristic profiles (Table S2), except for an exclusive expression of genes associated to KEGG pathways related to cancer after VILDA treatment (Table S2). In general, our findings underline a rather mild effect of VILDA compared to OA and for further analyses, we included both the exclusively and common but differentially expressed genes.

**Fig. 6.**
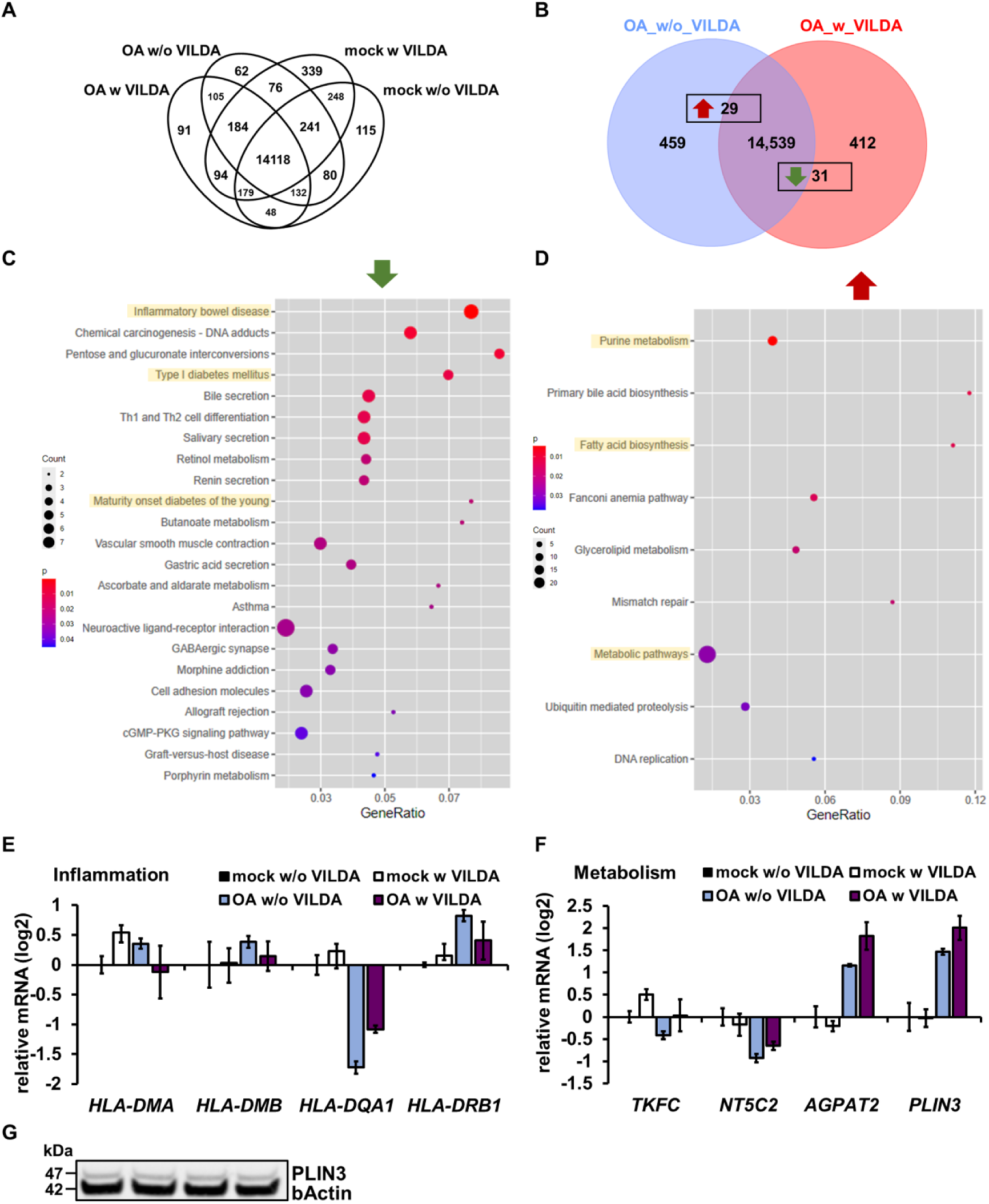
VILDA effects on the global transcription of steatotic Cntrl 1 HLCs. **(A)** Venn diagram of expressed genes upon indicated treatments. **(B)** Venn diagram of the gene expression of OA treated HLCs w and w/o VILDA. **(C,D)** KEGG-associated pathways of significantly down-**(C)** and up-**(D)** regulated genes upon OA w VILDA treatment (complete gene lists in supplementary table S4). **(E,F)** Gene expression of inflammation-associated **(E)** and metabolism-associated **(F)** genes upon OA w and w/o VILDA treatment. **(G)** Cropped WB of Perilipin-3 (PLIN3) upon OA w and w/o VILDA in comparison to housekeeping protein b-Actin. Uncropped full-length blots can be found in the supplementary Fig. S9.

Next, we wanted to know, whether VILDA influences gene expression upon OA treatment and found 29 genes significantly up- and 31 genes significantly downregulated upon VILDA treatment (Fig. 6B). KEGG-pathway analysis revealed genes belonging to the inflammatory bowel disease pathway, asthma and Type 1 diabetes mellitus (T1DM) associated pathways significantly downregulated (Fig. 6C and Fig. S7A). Common downregulated genes of these pathways encode members of the Human leukocyte antigen II (HLAII) family, namely HLA-DQA1 and HLA-DMB, which are known to be ectopically expressed on hepatocytes upon hepatitis(35–37). Although the expression changes were small, we confirmed upregulation of *HLA-DMA*, *HLA-DMB* and *HLA-DRB1* upon OA-treatment and a tendency towards downregulation upon OA w VILDA (Fig. 6E). Interestingly, in contrast to the RNA-seq we found *HLA-DQA1* downregulated upon OA and a slight increase of expression upon OA w VILDA treatment (Fig. 6E). These data might indicate a potential association between DPP4 and HLAs in steatosis. However, experiments with more cell lines and prolonged treatment would be necessary for confirmation.

Genes associated with metabolic pathways, such as purine metabolism and fatty acid biosynthesis pathways were upregulated in OA w VILDA versus OA w/o VILDA (Fig. 6D and Fig. S7A). Interestingly, we found triokinase and FMN cyclase (*TKFC*), and 5’-nucleotidase, cytosolic II (*NT5C2*), both members of the purine metabolism, downregulated upon OA w/o VILDA in comparison to mock w/o VILDA, while VILDA treatment induced an upregulation, which might indicate a restoration of the pathway (Fig. 6F). Considering fatty acid metabolism, we detected O-acyltransferase 2 (*AGPAT2*) upregulated upon OA w/o VILDA treatment in comparison to mock w/o VILDA which was even reinforced upon OA w VILDA (Fig. 6F). We saw the same trend in gene expression of perilipin-3 (*PLIN3*) however, we did not detect differential protein expression (Fig. 6F-G). In contrast, considering another member of the perilipin family, namely PLIN2, we detected the same gene expression pattern, but differential protein expression (Fig. S6B, Fig. 6F). PLIN2 protein levels were upregulated upon OA, while PLIN3 protein levels were stable throughout the conditions. A reason might be, that PLIN3 belongs to the exchangeable PLINs, while PLIN2 is a constitutive protein, upregulated upon OA-treatment and unstable in the absence of LDs(38). PLIN3 is stable in the cytoplasm independently of LDs but is recruited to the lipid fractions(38). Nevertheless, considering the mRNA expression, our findings might indicate that DPP4 inhibition upon OA restores the hampered cellular energy homeostasis, however further studies are necessary to pinpoint the underlying mechanism.

To test whether VILDA affects the expression of steatosis-associated genes, we analyzed the expression of genes belonging to the PPAR and gluconeogenesis pathways, which were both relevant in our previous studies, via heatmaps and detected clustering according to each condition (Fig. S10, Fig. S11). In addition, we evaluated the steatosis gene set from the earlier global analysis. We could confirm the same trend of gene expression for Cntrl 1 upon OA (Fig. S6A), while we could not detect a significant difference when comparing OA w/o and w VILDA.

To gain insights into the overall VILDA effect, independent of the steatosis condition, we performed KEGG-pathway analysis of the genes differentially expressed under mock conditions. We found 47 genes significantly upregulated and 44 genes significantly downregulated upon mock w and w/o VILDA treatment (Fig. S8A). KEGG-associated pathway analysis revealed, among others, downregulated genes associated with inflammatory response pathways under mock conditions (Fig. S8B). Upregulated genes were involved in KEGG-associated pathways such as enhanced metabolism and insulin secretion, indicating beneficial effects for the compromised insulin pathway, since insulin resistance pathway was upregulated upon OA (Fig. S8C). Altogether, our findings support the hypothesis that DPP4 is involved not only in metabolic regulation via purine and fatty acid metabolism but furthermore affects inflammatory-associated pathways.

## Discussion

### Oleic acid feeding induces clinically relevant phenotype of steatosis

In this study, we generated induced pluripotent stem cell (iPSC)-derived steatotic hepatocyte-like cells (HLCs) from four individuals to elucidate the potential hepatocyte-specific contribution to the progression of MAFLD. Although iPSC-derived HLCs in general are not fully mature, we and others could show that they have considerable metabolic activity which makes them a valuable system to model MAFLD, reflecting the diverse genetic backgrounds of the donors(7, 8, 39–41). By stimulating HLCs with OA, we were able to induce lipid droplet formation in all four cell lines, as already shown in previous studies(7, 8). Although two of the cell lines were derived from steatosis patients, there was no disease-specific effect w or w/o OA detectable. This is specifically interesting, since our previous findings indeed indicated steatosis related gene expression patterns(9). Nevertheless, cell line-specific differences in the amount and size of the lipid droplets were observed. This is in line with our previously published data and the high divergence in individual symptoms and progression of MAFLD which are well-known difficulties in the clinic(7). They are due to genetic variations like single nucleotide polymorphisms (SNPs), epigenetic alterations and other co-morbidities that are associated with the disease(40, 42). Furthermore, considerable variability is typical for iPSC-derived data due to the individual genetic background and differences in differentiation efficiencies. Indeed, we excluded the patient derived cell line Stea 2 due to ambiguous clustering in the first level analysis.

Global transcriptomic analyses revealed MAFLD-associated pathways such as metabolism-associated and immune-modulating pathways differentially regulated in our model(43–45). A heatmap analysis of genes from these pathways, revealed clustering according to the treatment and independent of the genetic background. This underlines the importance of the involved genes for the disease. Although our model comprises only hepatocytes, the detected upregulation of cytokine-cytokine receptor signaling, NF-kappa B and TNF signaling upon OA might indicate immune cell recruitment via inflammatory/ chemokine signaling. Recently, Yu et al demonstrated that hepatocyte-intrinsic changes contribute to the disease(44) while predisposition, environment, or other comorbidities might further regulate the pace of the progression towards fibrosis and cirrhosis. Comparing the gene expression profile detected in our HLCs with their single cell RNA-seq data from liver resections of NASH and HCC-patients, we found many pathways upregulated that are associated with a lower risk of NASH-HCC-transition, such as galactose catabolic processes, hexose metabolic processes, and glucose homeostasis.

### Active Dipeptidyl peptidase 4 (DPP4) is secreted upon OA treatment

DPP4 is a serine protease with catalytic activity for various substrates(12) and considered a hepatokine, upregulated in metabolic liver disease and driver of inflammation(13). Similar to observations made in HepG2 cells(46), we could not detect elevated DPP4 on both the mRNA or protein level. However, we found elevated secretion and a drastic increase in the catalytic activity of DPP4 upon OA, in line with previously observed elevation in NAFLD/NASH patients(47, 48). As DPP4 is involved in various chronic and cancerous diseases throughout the human body, hepatocyte-specific secretion upon late-steatosis might indicate inflammatory signaling and a risk for the development of other comorbidities like cardiovascular and renal diseases(49).

### VILDA interferes with DPP4 activity and regulates pathways related to inflammation and metabolism

As a first insight into the mechanism of action of DPP4, we inhibited its activity with VILDA-an FDA-approved T2DM medication. As a proof-of-principle, we found that VILDA interferes solely with DPP4 activity and not with the protein or mRNA level, as described in previous studies(50). In our study, as the first in this format, global transcriptome analysis revealed clustering according to the OA-/mock treatment, but not according to VILDA treatment. This confirmed our expectation that OA treatment induced greater transcriptome changes than VILDA treatment. Nevertheless, we detected exclusively expressed genes for all four conditions. Although KEGG-associated pathway analysis did not identify characteristic profiles, we noticed that genes involved in the development of cancer were upregulated upon VILDA independently of steatosis. VILDA-associated safety concerns have already been addressed extensively and no significant overall cancer-association was found(51, 52). Considering the typical long-term or even life-long medication of T2DM, this should nevertheless be monitored carefully.

Furthermore, we found inflammation associated pathways such as Type 1 diabetes mellitus (T1DM), inflammatory bowel disease and asthma downregulated upon DPP4 inhibition. *HLA-DQA1* and *HLA-DMB* are common genes involved in all of them, and their expression was differentially regulated upon steatosis and additional DPP4 inhibition. HLA class II proteins are typically expressed on the surface of antigen presenting cells, however ectopic expression in hepatocytes upon disease has been shown(35, 37). In line with this, we found a tendency of elevated gene expression of *HLA-DMB*, *HLA-DMA* and *HLA-DRB1* upon OA, which are associated with NASH, hepatitis and cirrhosis(53, 54), and all decreased upon DPP4 inhibition. Since DPP4 activity affects (auto)-immune related diseases in a complex manner(55) this might provide a possible clue towards its role for steatosis/MAFLD progression. However, further studies are necessary to elucidate the role of DPP4 in the steatosis model.

To understand the role of DPP4 for the interplay between metabolism and inflammation, it is essential to determine, whether DPP4 is causal or correlative for late-steatosis. DPP4 was shown to be epigenetically regulated(56, 57). Indeed, we also found a slight, albeit not significant demethylation upon OA treatment (not shown). This supports the speculation that early events of energy overload might change the methylation profiles of CpG islands in the DPP4 locus and enable DPP4 expression in hepatocytes at a rather early timepoint of disease progression.

The insulin resistance pathway was upregulated upon OA treatment, which matches with the well-known insulin resistance promoting effect of DPP4. In addition, genes involved in PPAR signaling and gluconeogenesis showed condition dependent gene expression patterns. Interestingly, we found genes involved in purine and fatty acid metabolism upregulated upon VILDA treatment. E.g. *AGPAT2,* which is involved in fatty acid metabolism, was upregulated upon DPP4 inhibition. Its deletion or mutation is associated with insulin resistance, diabetes and severe forms of metabolic syndrome in mice and humans(58–60). This could indicate a VILDA-mediated beneficial effect for the hampered metabolism due the energy overload during late-steatosis and underline the idea of a role for DPP4 in the interplay between metabolism and inflammation(16, 61).

### Conclusions

Taken together, we provide a human iPSC-derived model focusing on the hepatocyte-specific contribution to progression of steatosis. We could link DPP4 activity to the steatosis phenotype and show that its inhibition with Vildagliptin has effects on metabolism- and inflammation-associated gene expression during steatosis. Since we have performed global transcriptome analyses of the effects of VILDA with only one cell line, these can only provide first insights into possible effects, and more in depth analyses are needed. In the future, human DPP4-knockout HLCs, embedded in a multicellular liver model, could increase our understanding of the mechanisms of DPP4 through health and disease and help to further elucidate the interplay between the distinct cell types of the liver.

## Abbreviations

ACE2: Angiotensin converting enzyme 2
AFP: Alpha-Fetoprotein
AGPAT2: O-acyltransferase 2
ALB: Albumin
AMPK: Adenosine monophosphate-activated protein kinase
ANPEP: Alanyl Aminopeptidase
CCL4: C-C motif chemokine ligand 4
CD26: Cluster of differentiation 26
Cntrl: Control
CPT1A: Carnitine palmitoyltransferase I
CYP: Cytochrome P450
DE: Definitive endoderm
Dex: Dexamethasone
DPP4: Dipeptidyl peptidase 4
E-CAD: E Cadherin
ELISA: Enzyme-linked immunosorbent assay
FABP1: fatty acid binding protein 1
FDA: Food and drug administration (U.S.)
GOs: Gene ontologies
GSK-3: Glycogen synthase kinase 3
HE: Hepatic endoderm
HGF: Hepatocyte growth factor
HLAII: Human leukocyte antigen II
HLCs: Hepatocyte like cells
HNF4a: Hepatocyte nuclear factor 4alpha
ICC: Immunocytochemistry
IGFBP1: Insulin Like Growth Factor Binding Protein 1
igG: Immunoglobulin G
IL1B: Interleukin 1beta
iPSCs: induced pluripotent stem cells
KEGG: Kyoto Encyclopedia of Genes and Genomes
LD: Lipid droplet
MAFLD: Metabolic dysfunction-associated fatty liver disease
MASLD: Metabolic dysfunction-associated steatotic liver disease
NAFLD: Non-alcoholic fatty liver disease
NASH: Non-alcoholic steatohepatitis
NT5C2: 5’-nucleotidase, cytosolic II
OA: Oleic acid
OCT4: octamer-binding transcription factor 4
OSM: Oncostatin M
P/S: Penicillin/Streptomycin
PCK1: Phosphoenolpyruvate Carboxykinase 1
PLIN2: Perilipin-2
PLIN3: Perilipin-3
PPAR: Peroxisome proliferator activated-receptor
qRT-PCR: Quantitative reverse transcription PCR
R.L.U.: Relative light units
RNA-seq: RNA sequencing
ROS: Reactive oxygen species
RPLP0: Ribosomal Protein Lateral Stalk Subunit P0
SNPs: single nucleotide polymorphisms
SOX: SRY-related HMG-box genes
Stea: Steatosis
T1DM: Type 1 diabetes mellitus
T2DM: Type 2 diabetes mellitus
TBS-T: Tris-buffered saline with Tween20
TKFC: triokinase and FMN cyclase
TNF: Tumor necrosis factor
VCAM1: Vascular cell adhesion molecule 1
VILDA: Vildagliptin

## Declarations

### Ethics approval and consent to participate

The use of iPS cell line Cntrl 1 was approved by the ethics committee of the medical faculty of Heinrich Heine University Düsseldorf under ethical approval number 5704 on the 01.03.2017. The title of the approved project is „Isolierung, Charakterisierung und Reprogrammierung von Stammzellen aus dem Urin”. The use of iPS cell lines Cntrl 2, Stea 1 and Stea 2 was approved by the ethics committee of the medical faculty of Heinrich Heine University Düsseldorf under ethical approval number 5013 on the 09.07.2015. The title of the approved project is „Nutzung humaner embryonaler Stammzellen, fetaler humaner Stammzellen (MSC) sowie induzierter pluripotenter Stammzellen zur Erforschung von Lebererkrankungen (NAFLD) und Hirnerkrankungen (Alzheimer)“. The patients or their guardian(s)/legally authorized representative(s)/next of kin provided written informed consent for participation in the study and/or the use of samples.

### Availability of data and materials

The data that support the findings of this study are available from the corresponding author upon reasonable request. NGS data will be provided on the GEO-server and/or on upon reasonable request upon acceptance. Additional files of produced sequencing data is provided in supplementary table S3 and S4.

## Acknowledgments

C.L., N.G. and J.R. are funded by the Else Kröner-Fresenius-Stiftung – 2020_EKEA.64. J.A. acknowledges the medical faculty of Heinrich-Heine University Düsseldorf for part funding of this project. The authors declare that they have not use AI-generated work in this manuscript.

## Authors contribution

CL: Conceptualization, Data curation, Investigation, Methodology, Visualization, Writing–original draft, Writing–review and editing. WW: Data curation, Visualization, Writing–original draft, Writing–review and editing JR: Investigation, Methodology, JA: Conceptualization, Project administration, Resources, Supervision, Writing–original draft, Writing–review and editing. NG: Conceptualization, Formal Analysis, Funding acquisition, Investigation, Methodology, Project administration, Resources, Supervision, Visualization, Writing–original draft, Writing–review and editing

## Authors information

Not applicable

