## Supplementary material for "DPP4 inhibition affects metabolism and inflammation associated pathways in hiPSC-derived steatotic HLCs"

**Supplementary Methods**

*Next generation sequencing and analysis of deep sequencing data*

The HISAT2 (version 2.1.0) software(1) was employed to align the fastq files received from the BMFZ core facility against the GRCh38 genome. Parameter optimizations(2) were applied resulting in the command:

hisat2 -p 7 -N 1 -L 20 -i S,1,0.5 -D 25 -R 5 --mp 1,0 --sp 3,0 -x hisatindex/grch38_r109 -U input.fastq.gz -S output.sam

Via SAMtools software(3) the resulting BAM files were sorted by coordinates. Read counts per gene obtained with the subread (1.6.1) featurecounts software(4) using the ENSEMBL annotation file Homo_sapiens.GRCh38.109.gtf and parameters –t exon –g gene_id. Within the R/Bioconductor environment data was normalized with the voom(5) algorithm from the limma package(6) filtering genes, which were expressed with CPM (counts per million) > 1 in at least one sample. Venn diagrams were drawn with the VennDiagram package(7) based on genes considered expressed when there were more than 5 reads. Differential expression was determined by a p-value < 0.05 from the limma test and a fold change greater than 1.5 for genes expressed at least in one condition. The False-Discovery-Rate (FDR) was calculated by the method of Storey et al. implemented in the Bioconductor package qvalue (8) The complete correlation table and gene lists for OA/mock experiments can be found in the supplementary table 1. The complete correlation table and gene lists for OA w and w/o VILDA experiments can be found in the supplementary table 2.

*Microscopic imaging data*

**Make and model of microscope:** Zeiss, LSM 700 microscope

**Type, magnification, and numerical aperture of the objective lenses:** LD Plan-Neofluar 20x/0.4 and Plan-Apochromat 40x/1.4 Oil DIC (UV)VIS-IR

**Temperature:** 20-22 °C

**Imaging medium:** Phosphate-buffered saline (PBS) w/o magnesium and calcium, 1x Fluoromount-G W DAPI (Biozol, Cat. Number: SBA-0100-20)

**Fluorochromes:** Alexa Fluor 488 Dye, Alexa Fluor 594 Dye, Alexa Fluor 555 Dye, Alexa Fluor 647 Dye, Hoechst 33342

**Camera make and model:** Zeiss, AxioCam MRM

**Acquisition software:** Zeiss**,** ZEN2012 (blue edition), Version 6.1.7601 for fluorescence microscopy and ZEN2011 SP3 (black edition), Version 8,1,6,484 for confocal microscopy. Images were processed using ZEN software, Version 3.10.103.00000,

### Supplementary Figures

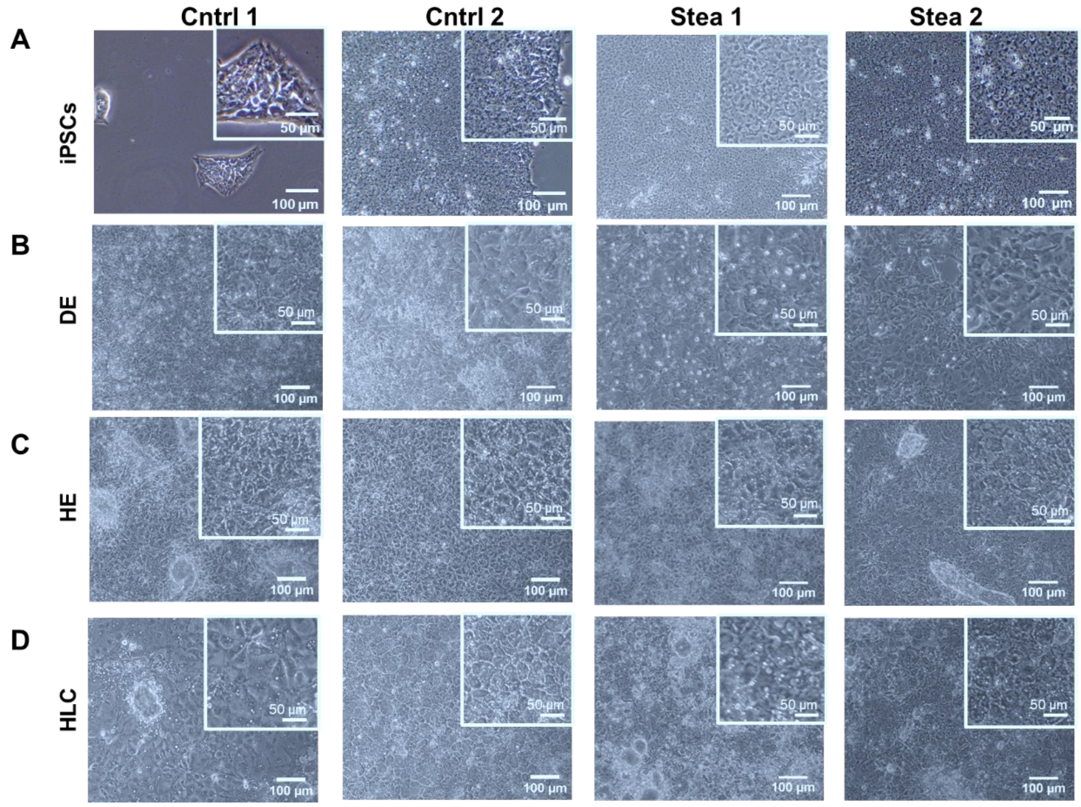

*Fig. S1 Cell morphology during the differentiation stages of cell lines Cntrl 1, Cntrl 2, Stea 1 and Stea 2.* Scale bars represent 100 µm and 50 µm in the zoom-in. **(A)** induced pluripotent stem cells (iPSCs). **(B)** Definitive endoderm (DE). **(C)** Hepatic endoderm (HE). **(D)** Hepatocyte-like cells (HLCs).

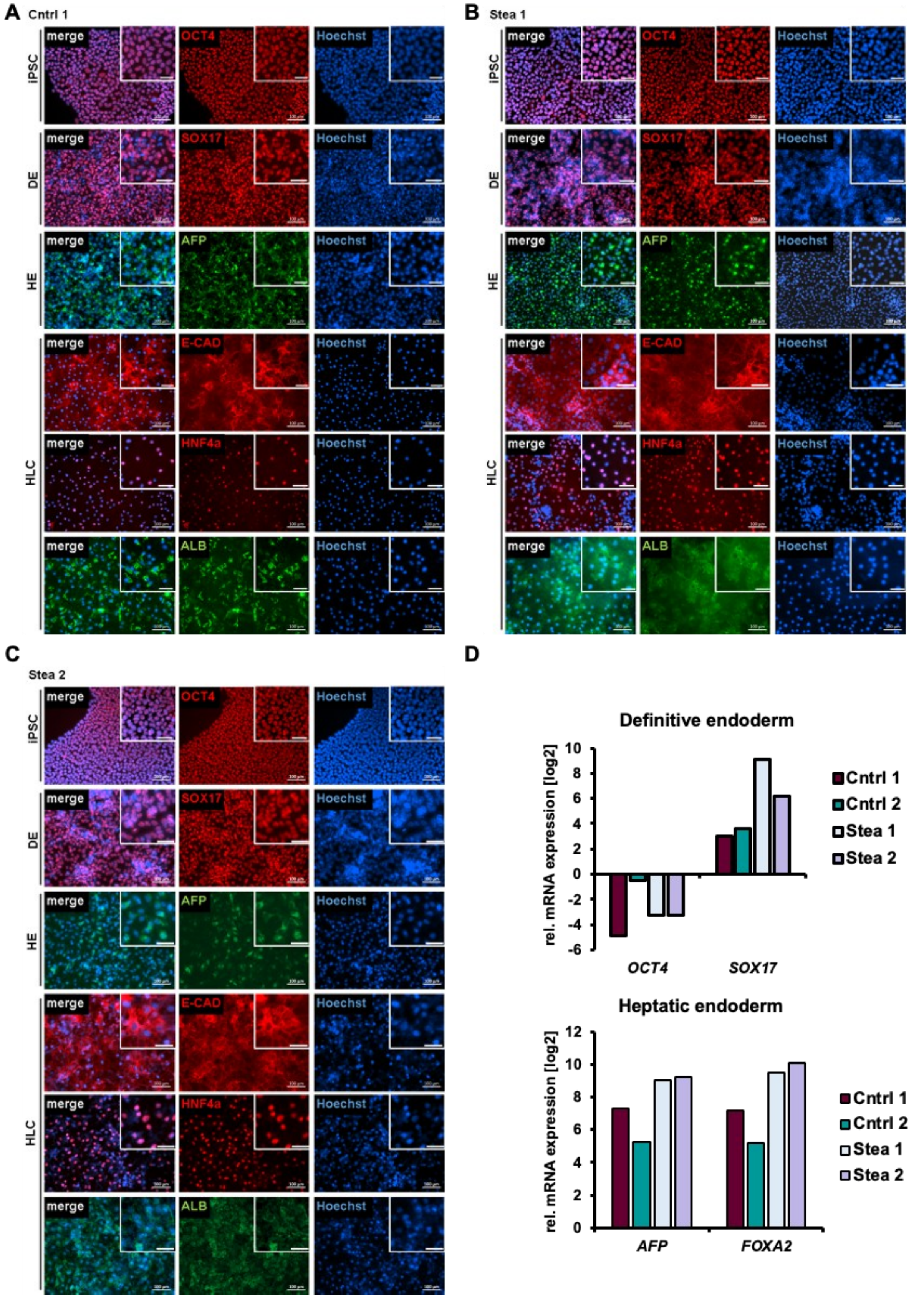

*Fig. S2 Characterization of HLC differentiation.* **(A-C)** Representative immunocytochemistry of cell line Cntrl 1, Stea 1 and Stea 2 of differentiation stages showing respective markers OCT4 in induced pluripotent stem cells (iPSCs), SOX17 in Definitive Endoderm (DE) and AFP in Hepatic Endoderm (HE). For hepatocyte-like cells (HLCs), the epithelial marker E-CAD, HNF4alpha and ALB are shown. **(D)** Gene expression of HLCs derived from four cell lines. Shown are means of three technical replicates (n=1) of *OCT4* and *SOX17* in DE and *AFP* and *FOXA2* in HE in comparison to iPSC-stage (n=1).

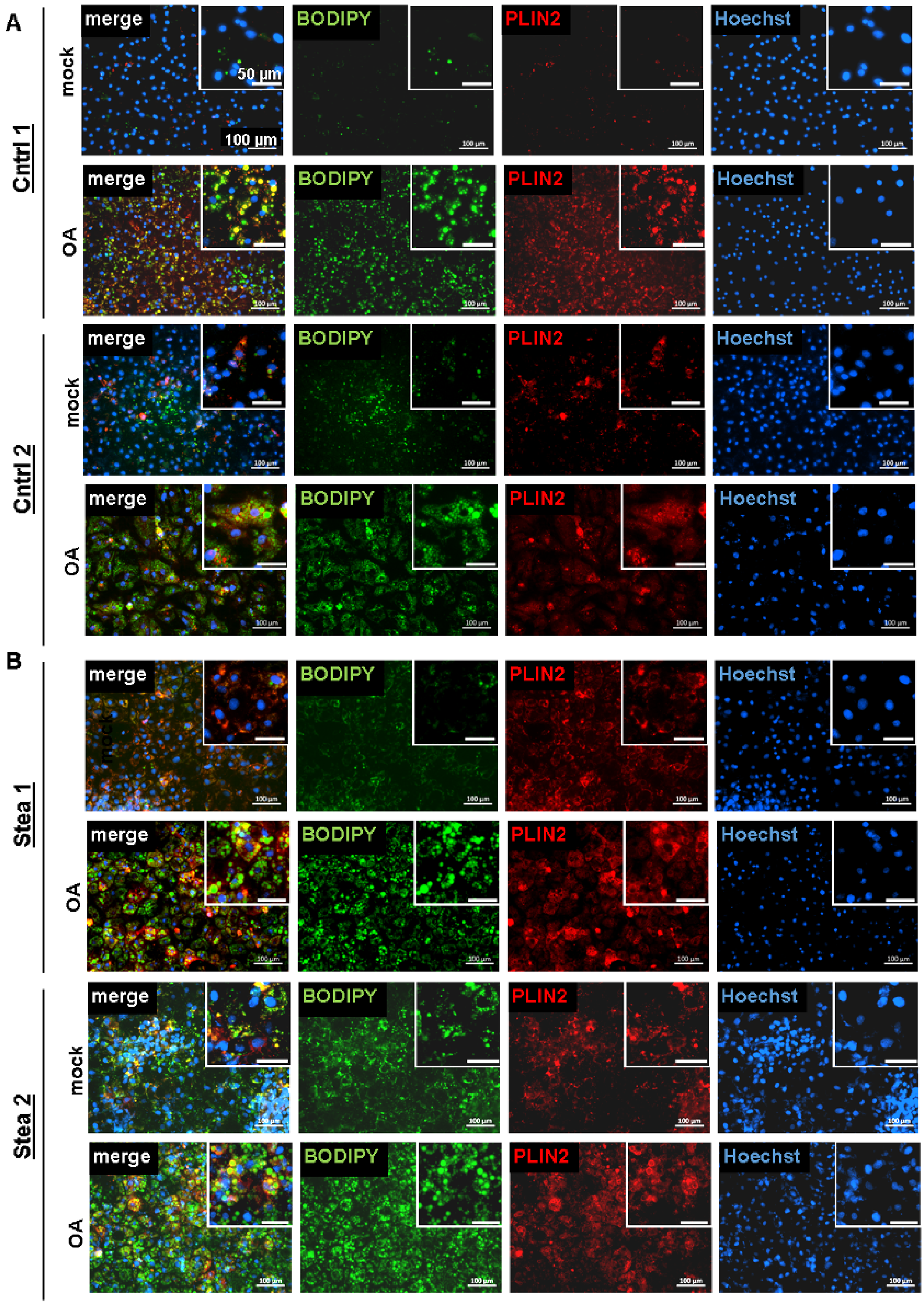

*Fig. S3 OA-induction of Lipid droplets.* **(A/B)** Representative immunofluorescence and BODIPY493/503 staining of HLCs of Cntrl 1, Cntrl 2 Stea 1 and Stea 2 HLCs treated with 400 µM OA (OA) and respective control (mock) for 7 days. PLIN2 is shown in red, fatty acids are stained in green, scale bars represent 100 µm in the original image and 50 µm in the zoom-in.

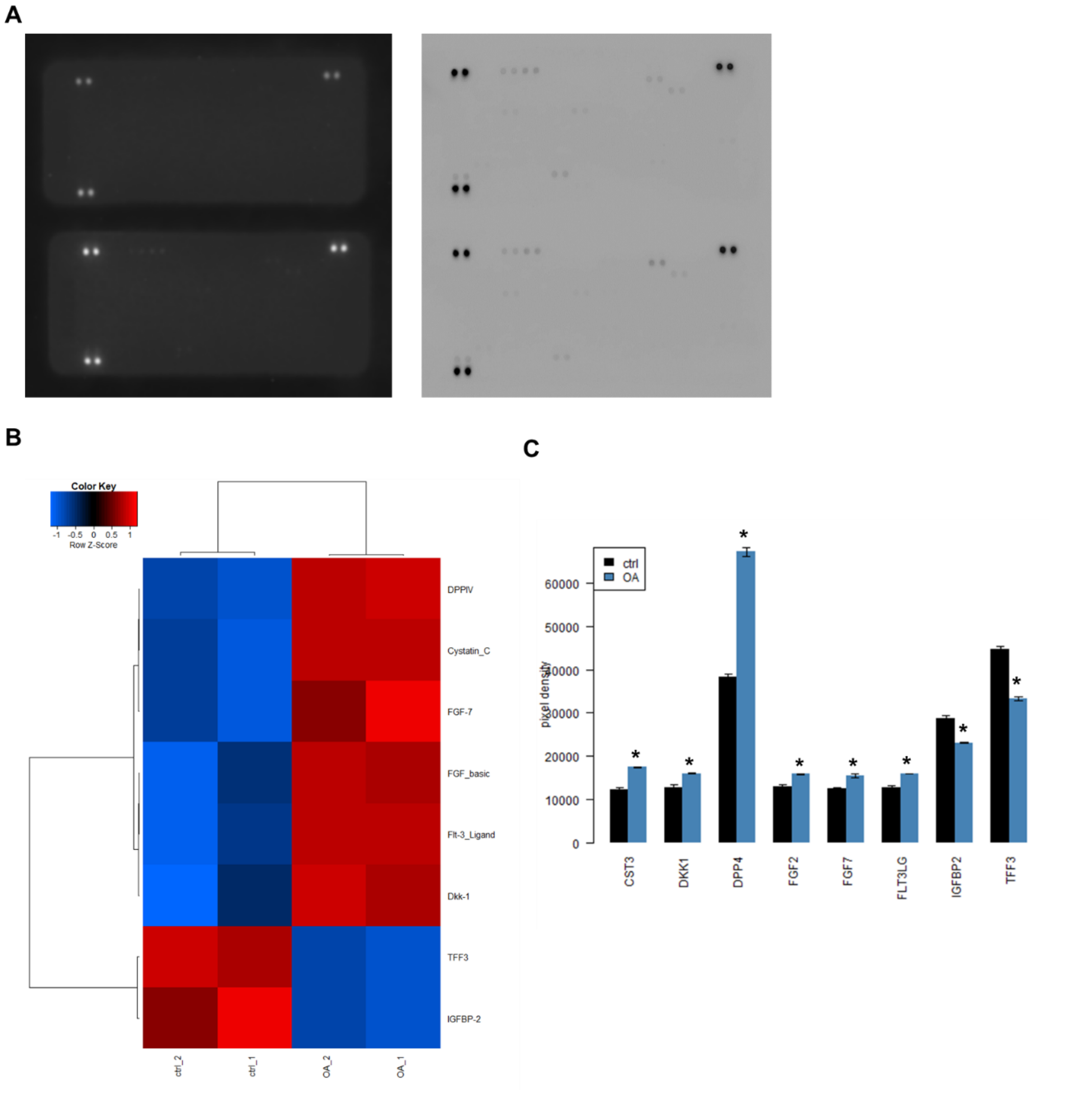

*Fig. S4 Released proteins after OA treatment.* Cntrl 1 HLCs were treated with OA for 7 days and the supernatant was analyzed for secreted proteins. **(A)** Array membranes, incubated with pooled supernatant of three biological replicates from mock (upper)- or OA (lower) treated Cntrl 1 HLCs. **(B)** Heatmap indicating significantly regulated proteins after analysis of the captured proteins on the membranes in technical duplicates. **(C)** Histogram of the detected chemiluminescence signal in pixel density for captured proteins under mock (ctrl) and OA treatment (* p-value < 0.05).

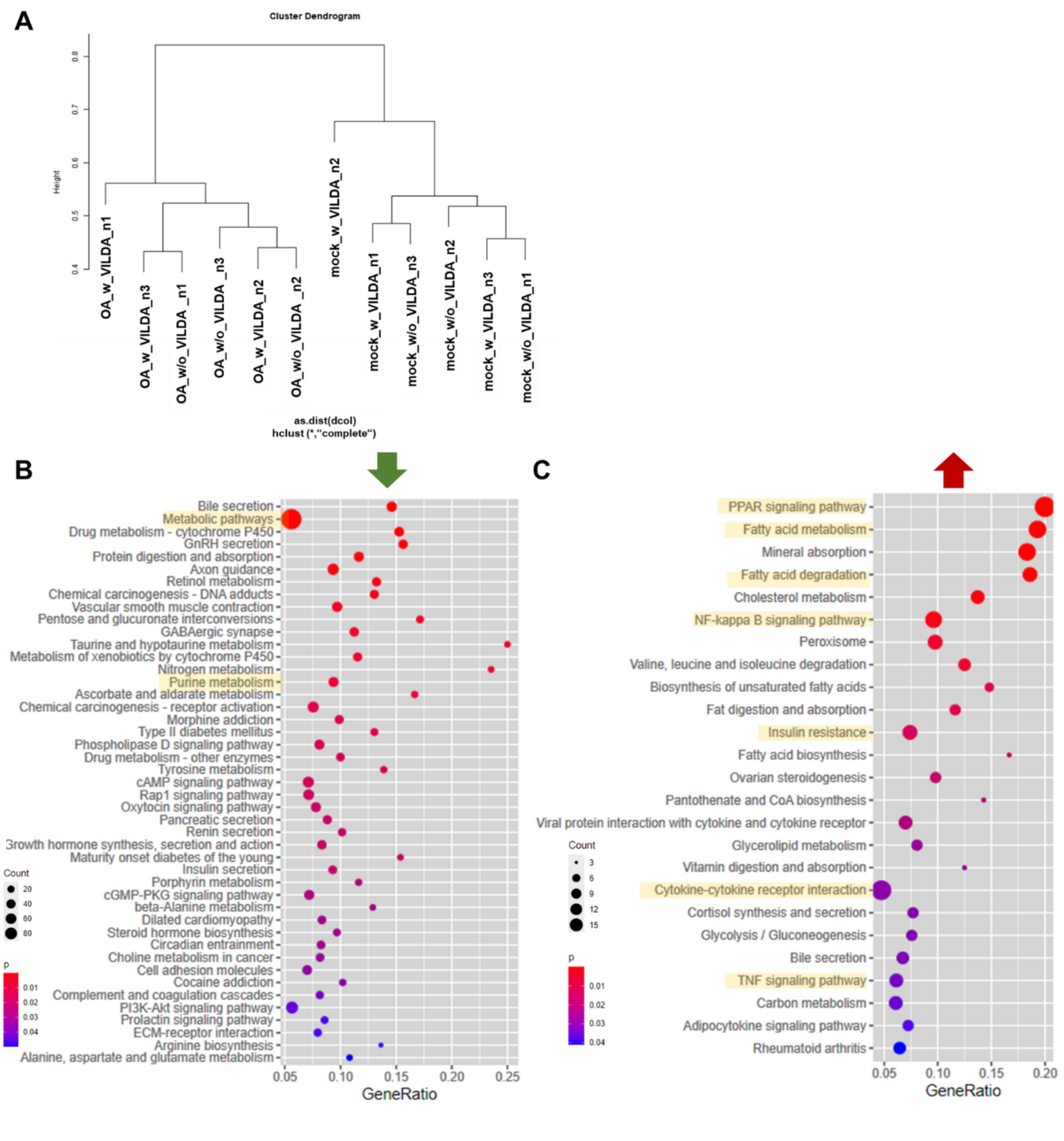

*Fig. S5 Global transcriptome analysis of HLCs treated with OA with and without VILDA.* **(A)** Hierarchical cluster dendrogram of global transcriptomic changes upon OA (mock or OA) with and without VILDA (w or w/o VILDA) treatment of HLCs derived from cell lines Cntrl 1 in three biological replicates (n=3). **(B)** KEGG-associated pathway analysis of significantly downregulated genes upon OA w/o VILDA in comparison to mock w/o VILDA treatment. **(C)** KEGG-associated pathway analysis of significantly upregulated genes upon OA w/o VILDA in comparison to mock w/o VILDA treatment.

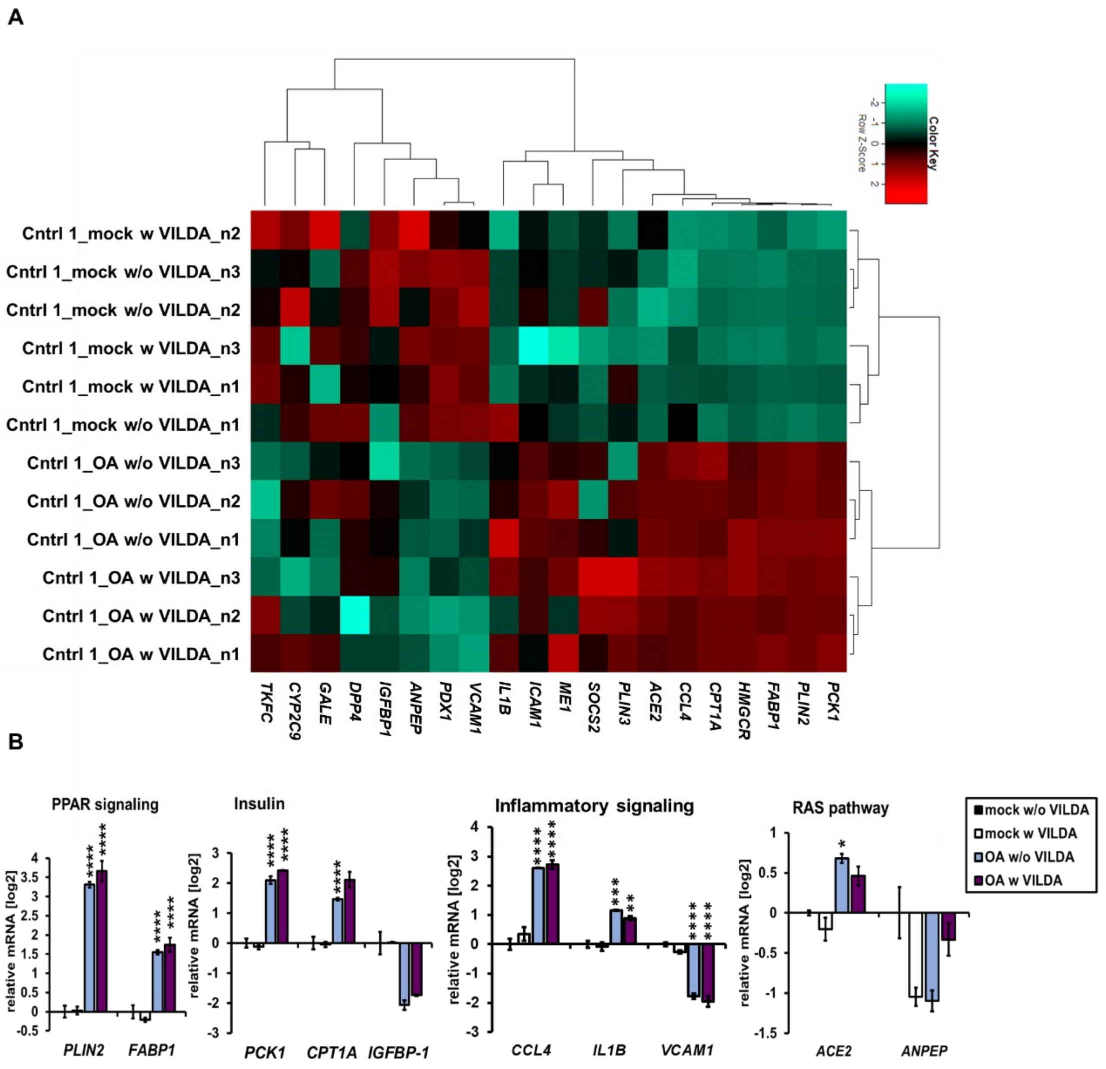

*Fig. S6 Gene expression of steatosis markers in HLCs treated with OA w and w/o VILDA.* **(A)** Heatmap analysis of steatosis associated genes. **(B)** Gene expression of *PLIN2*, *FABP1*, *PCK1*, *CPT1A*, *IGFBP1*, *CCL4*, *IL1B*, *VCAM1, ACE2*and *ANPEP*, shown as means of three biological replicates (n=3 ±SD), normalized to mock w/o VILDA. Ordinary two-way ANOVA, followed by Tukey`s multiple comparison test was performed to calculate significances (*p < 0.05, **p < 0.01, ***p < 0.001 in comparison to mock w/o VILDA).

A

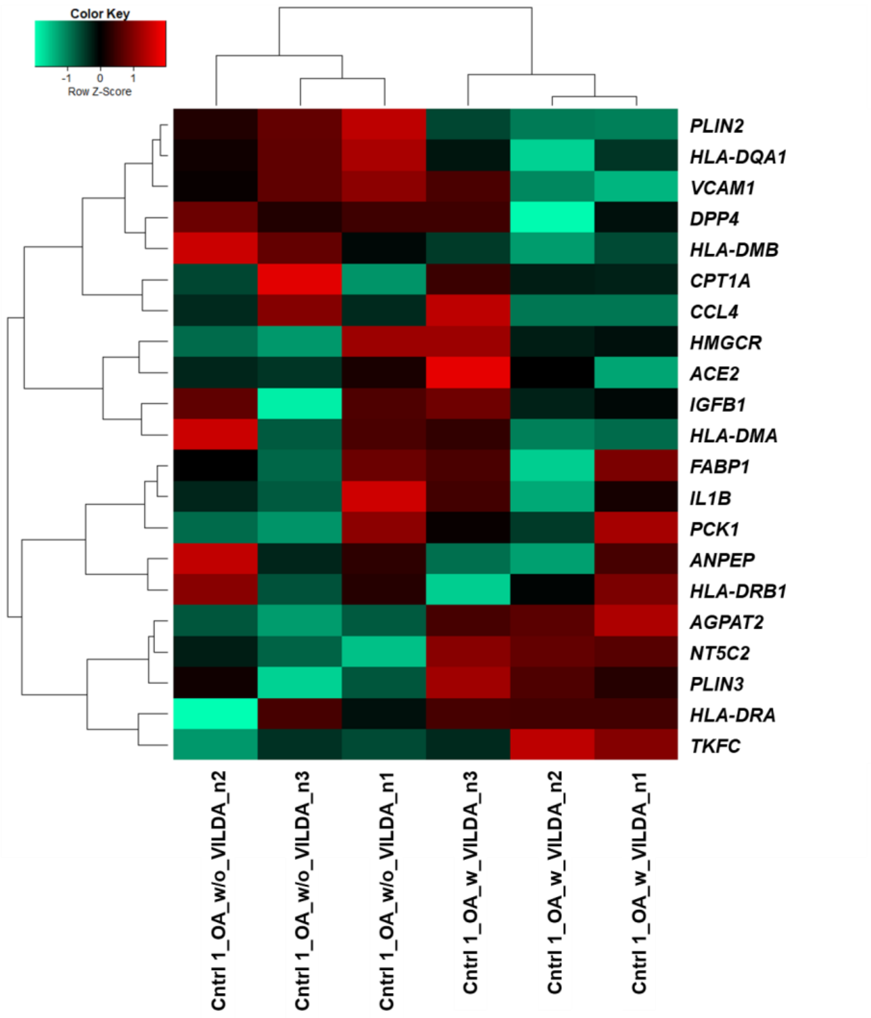

*Fig. S7 Pearson's correlation heatmap analysis upon OA w and w/o VILDA.* **(A)** Pearson's correlation heatmap analysis of genes involved in KEGG-associated pathways of purine metabolism (*TKFC*, *NT5C2*), fatty acid metabolism (*FABP1*, *PLIN3*, *AGPAT2*), inflammatory bowel disease, asthma and maturity onset diabetes of the young (*HLA-DQA1*, *HLA-DMB*).

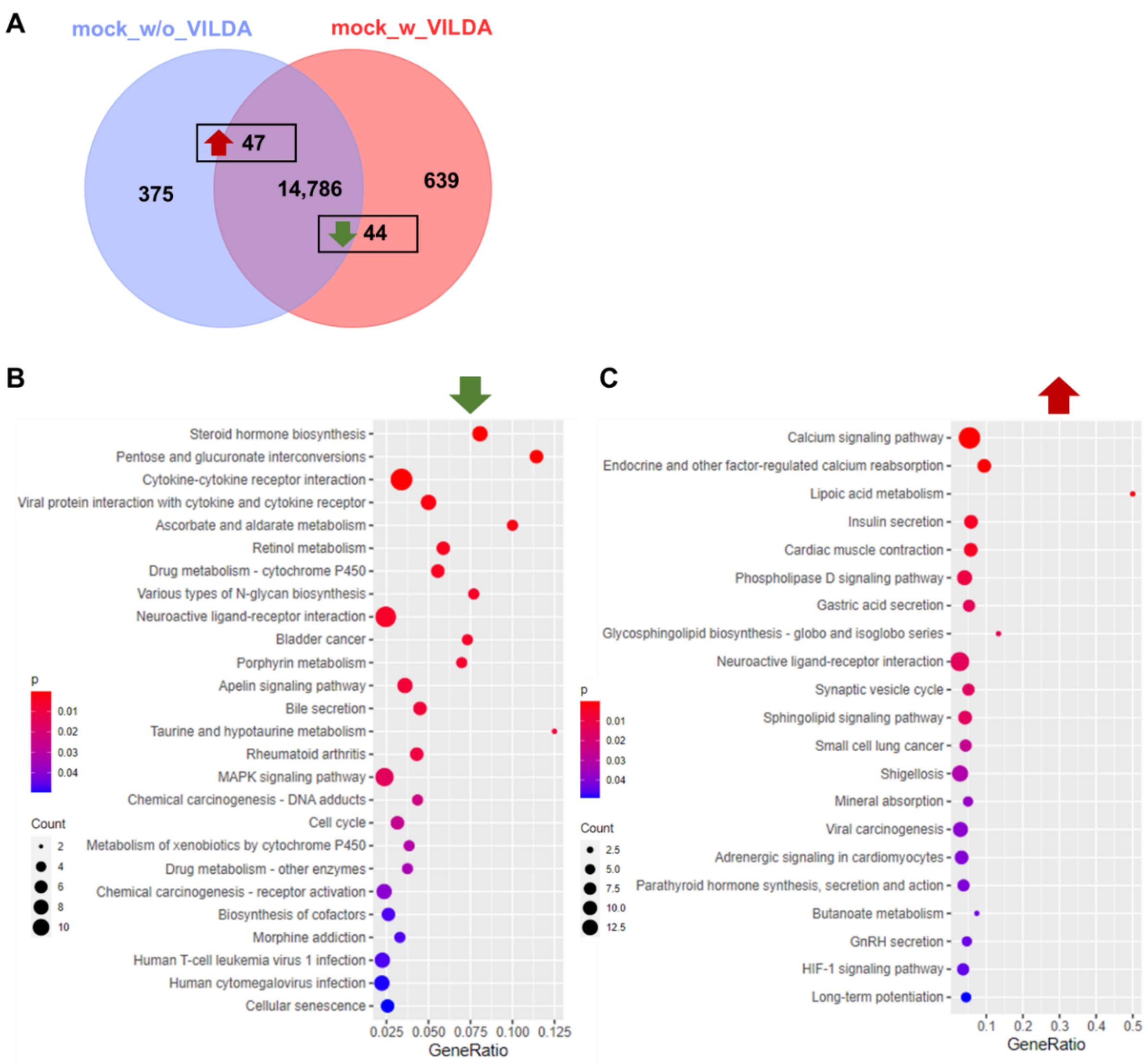

*Fig. S8 Global transcriptome analysis of mock-treated HLCs with and without VILDA.*
**(A)** Venn diagram of expressed genes upon mock treatment, indicating 14,786 commonly expressed genes. 375 and 639 genes were exclusively expressed w/o or w VILDA, respectively. Among the exclusive and common gene sets, 47 genes were significantly up-, while 44 genes were significantly downregulated upon VILDA. **(B)** KEGG-associated pathway analysis of downregulated gene expression in HLCs treated with mock w and w/o VILDA. **(C)** KEGG-associated pathway analysis of upregulated gene expression in HLCs treated with mock w and w/o VILDA.

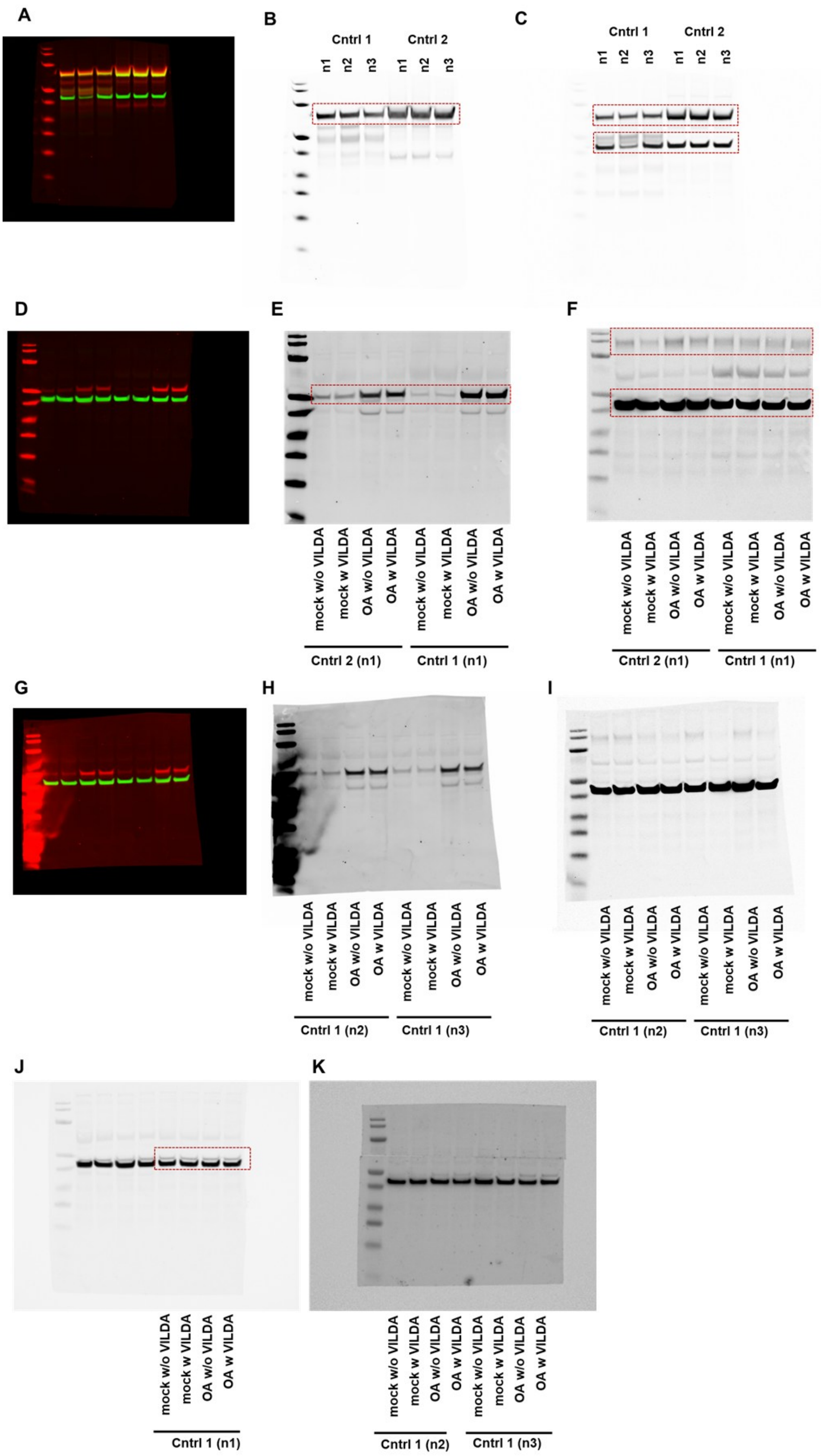

*Fig. S9 Uncropped full-length western blot membranes.*

**(A-C)** WB analysis of HLC differentiation, Fig. 1C. Peqlab peqGOLD Protein marker IV (VWR) was used to estimate the molecular weight of proteins. Bands (from bottom to top) represent proteins of the following weight in kDa: 10, 15, 25, 35, 40, 55, 100, 130, 170. **(A)** composite membrane detected in 680RD and 800RD channels. **(B)** AFP staining (1:2000 overnight, at 4 °C) and detection with IRDye 680RD goat anti-Rabbit (Licor) 1:10000, 1 h at RT. **(C)** ALB (1:5000 overnight, 4 °C) and bActin (1:5000, 4 °C) staining, detected with IRDye 800RD goat anti-Mouse (Licor) (1:10000, 1 h, RT). Dotted lines indicate the cropping. **(D-F)** Uncropped WB membrane for Fig. 5B: Analysis of HLCs treated with OA w and w/o VILDA n1. **(D)** Composite membrane detected in 680RD and 800RD channels. **(E)** PLIN2 staining (1:2000 overnight, at 4 °C) and detection with IRDye 680RD goat anti-Rabbit (Licor) 1:10000, 1 h at RT. **(F)** DPP4 (1:5000 overnight, 4 °C) and bActin (1:5000, 4 °C) staining, detected with IRDye 800RD goat anti-Mouse (Licor) (1:10000, 1 h, RT). Dotted lines indicate the cropping. **(G-I)** Full-length WB membrane for Fig. 5C: Analysis of Cntrl 1 HLCs treated with OA w and w/o VILDA n2 and n3. **(G)** Composite membrane detected in 680RD and 800RD channels. **(H)** PLIN2 staining (1:2000 overnight, at 4 °C) and detection with IRDye 680RD goat anti-Rabbit (Licor) 1:10000, 1 h at RT. **(I)** DPP4 (1:5000 overnight, 4 °C) and bActin (1:5000, 4 °C) staining, detected with IRDye 800RD goat anti-Mouse (Licor) (1:10000, 1 h, RT). **(J-K):** Uncropped WB membrane for Fig. 6G: Analysis of HLCs treated with OA w and w/o VILDA. PLIN3 Staining (1:5000 overnight, at 4 °C) and detection with IRDye 800RD goat anti-Mouse (Licor) (1:10000, 1 h, RT).

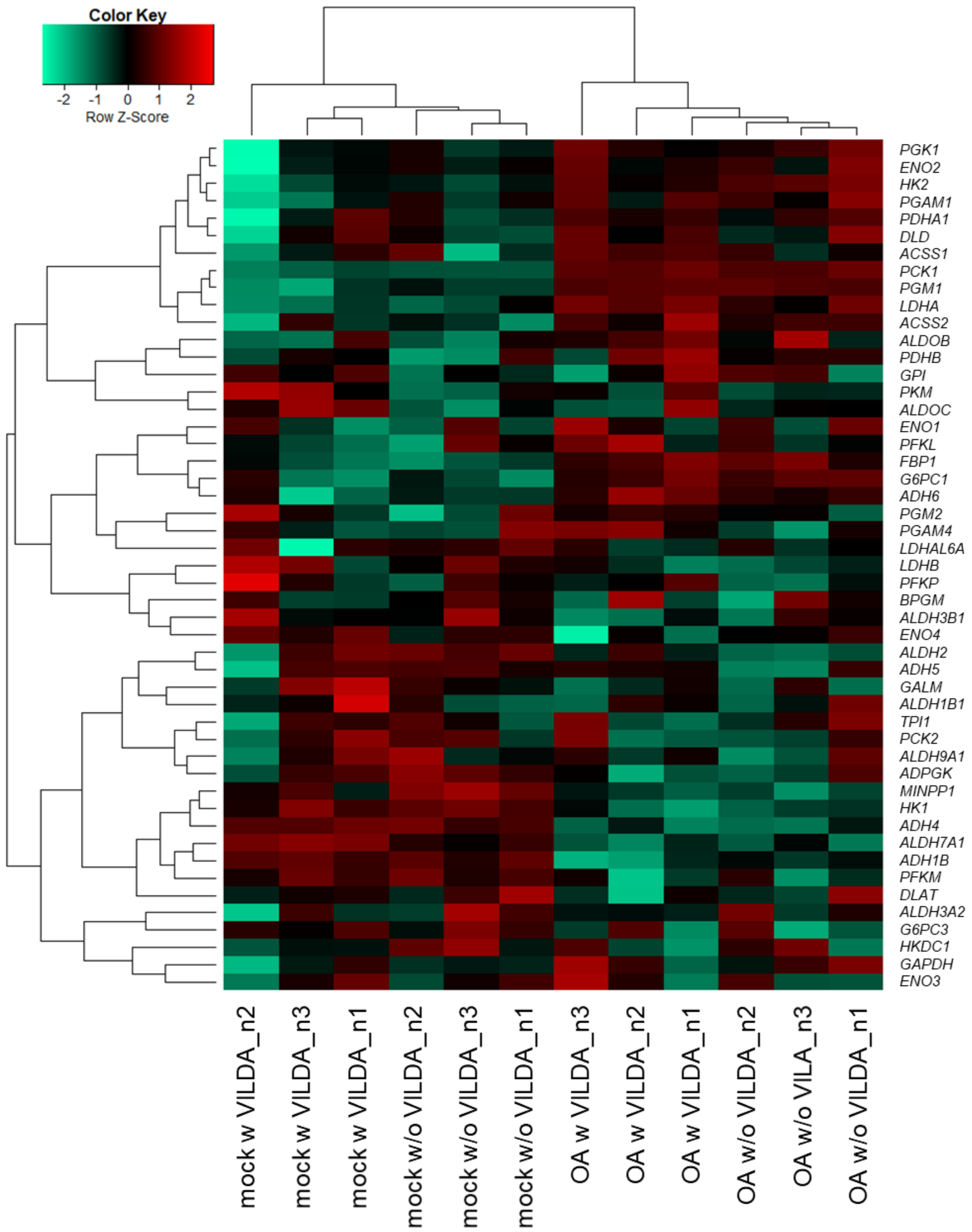

*Fig. S10 Person's correlation heatmap analysis of genes of the gluconeogenesis pathway.*

Pearson's correlation heatmap analysis of genes involved in KEGG-associated

pathways of gluconeogenesis.

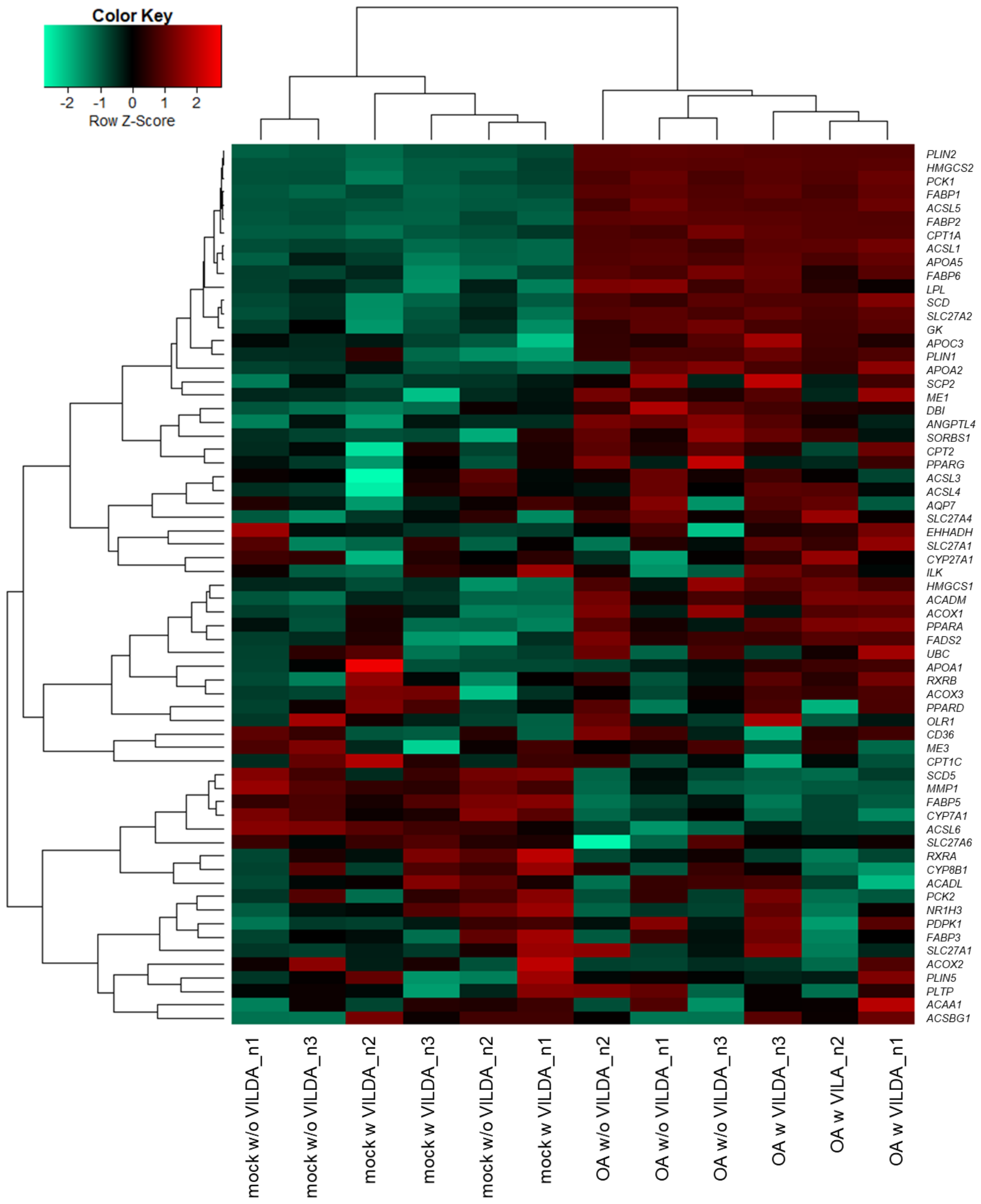

*Fig. S11 Person's correlation heatmap analysis of genes of the PPAR signaling pathway.*

Pearson's correlation heatmap analysis of genes involved in KEGG-associated pathway of PPAR signaling.

### Supplementary Tables

*Table S1: List of Primers*

| **Primers** | | |
|---|---|---|
| **Gene** | **Sequence** | |
| * | **Forward 5`-3`** | **Reverse 5`-3`** |
| *AFP* | AGCAGCTTGGTGGTGGATGA | CCTGAGCTTGGCACAGATCCT |
| *AGPAT2* | GGGGCGTCTTCTTCATCA | TTGAGGTTCTCCCTGACCAT |
| *Albumin* | AGCTGTTATGGATGATTTCGCAG | CCTCGGCAAAGCAGGTCTC |
| *ANPEP* | TGAGCTGTTTGACGCCATCT | GCCCTGCTTGAATACGTCCT |
| *CCL4* | GCTAGTAGCTGCCTTCTGCT | CCACAAAGTTGCCAGGAAGC |
| *CPT1A* | CCTACCACGGGTGGATGTTC | CAACATGGGTTTTCGGCCTG |
| *CYP2D6* | TTCCTGCCTTTCTCAGCAGG | GCACAAAGCTCATAGGGGGA |
| *CYP3A4* | GTGACTTTGCCCATTGTTTAGAAAG | CAGGCGTGAGCCACTGTG |
| *DPP4* | GTTCTTCTGGGACTGCTGGG | GCTGTAGCATCATCTGTGCCT |
| *FABP1* | ATCGTGCAGAATGGGAAGCA | CCCCTGTCATTGTCTCCAGC |
| *FOXA2* | TTCAGGCCCGGCTAACTCTG | CCTTGCGTCTCTGCAACACC |
| *HLA-DMA* | GGGTTTCCTATCGCTGAAGTG | CCAATAGGCAATTGCTGTGTA |
| *IGFBP1* | ACCATCACTTGCCCAGAGTT | AGGAGCAGTACCAGCCAGAC |
| *IL1B* | TGTACCTGTCCTGCGTGTTG | ACTGGGCAGACTCAAATTCCA |
| *OCT4* | AGTTTGTGCCAGGGTTTTTG | ACTTCACCTTCCCTCCAACC |
| *PCK1* | GGGAGTCTCCGGAAGGTGTT | CATGGCAAAGGGGTCATGC |
| *PLIN2* | GCTGAGCACATTGAGTCACG | TGGTACACCTTGGATGTTGG |
| *RPLP0* | TCGACAATGGCAGCATCTAC | ATCCGTCTCCACAGACAAGG |
| *sHLA-DMB* | CGGCCACCATCTGTGCAAGT | CCAGTCCCGAACGATGGGCT |
| *sHLA-DQA1* | GAAGGAGACTGCCTGGCG | CATGATGTTCAAGTTGTGTTTTGC |
| *sHLA-DRB1* | ACCCAAGCGTGACAAGCCCT | CCCGACTCCACTCAGCATCTTG |
| *SOX17* | ACGTGTACTACGGCGCGATG | CTGGTGCTGGTGCTGGTGTT |
| *VCAM1* | CGAACCCAAACAAAGGCAGAGTA | GAGGAAGGGCTGACCAAGACG |

*Table S2: List of Antibodies*

| Use: | | | | Western Blot | | Immunocytochemistry | |
|---|---|---|---|---|---|---|---|
| **Antibody** | **Host species** | **Manufacturer** | **Catalog number** | **Dilution** | **Diluent/ Blocking** | **Dilution** | **Diluent/Blocking** |
| Albumin (ALB) | mouse | Sigma | A6684-.2ml | 1:5000 | 5 % Milk TBS-T | 1:100 | 10 % goat serum |
| Alpha-fetoprotein (AFP) | rabbit | Sigma | HPA023600 | 1:2000 | 5 % Milk TBS-T | 1:300 | 3 % BSA |
| Angiotensin converting enzyme 2 (ACE2) | goat | R&D Systems | AF933 | 1:5000 | 5 % Milk TBS-T | NA | NA |
| beta-Actin | mouse | Cell Signaling Technologies | 3700S | 1:5000 | 5 % Milk TBS-T | NA | NA |
| Dipeptidyl peptidase 4 (DPP4) | mouse | Proteintech | 68383-1-Ig | 1:5000 | 5 % Milk TBS-T | 1:100 | 3 % BSA |
| E-Cadherin (ECAD) | goat | Cell Signaling Technologies | 3195 | NA | NA | 1:100 | 3 % BSA |
| Hepatocyte nuclear factor 4alpha (HNF4a) | rabbit | Abcam | 92378 | NA | NA | 1:250 | 3 % BSA |
| Octamer-binding transcription factor 4 (OCT4) | rabbit | Cell Signaling technologies | 2840S | NA | NA | 1:400 | 3 % BSA |
| Perilipin-2 (PLIN2) | rabbit | Proteintech | 15294-1-AP | 1:2000 | 5 % Milk TBS-T | 1:200 | 3 % BSA |
| Perilipin-3/TP47 (PLIN3) | mouse | Proteintech | 66523-1-Ig | 1:5000 | 5 % Milk TBS-T | NA | NA |
| Sry box transcription factor 17 (SOX17) | goat | R&D Systems | AF1924 | NA | NA | 1:50 | 3 % BSA |
| IRDye680RD Goat anti-Rabbit | goat | Licor | 925-68070 | 1:10,000 | 5 % Milk TBS-T | NA | NA |
| IRDye800RD Goat-anti-Mouse | goat | Licor | 925-68071 | 1:10,000 | 5 % Milk TBS-T | NA | NA |
| Alexa 594 gt anti ms IgG (H+L) | goat | Life technologies | A10521 | NA | NA | 1:500 | 3 % BSA |
| Alexa647 gt anti-Rabbit IgG (H+L) | goat | Life technologies | A32733 | NA | NA | 1:500 | 3 % BSA |
| Alexa488 gt anti-Rabbit IgG (H+L | goat | Life technologies | A11008 | NA | NA | 1:500 | 3 % BSA |
| Alexa555 dk anti-Goat IgG | donkey | Life technologies | A32816 | NA | NA | 1:500 | 3 % BSA |
